# A floristic survey of Fair Isle

**DOI:** 10.1101/164566

**Authors:** Camila V. Quinteros Peñafiel, Nick J. Riddiford, Alex D. Twyford

**Affiliations:** Royal Botanic Garden Edinburgh, Scotland, United Kingdom |; Schoolton, Fair Isle, Shetland ZE2 9JU |; Institute of Evolutionary Biology, The University of Edinburgh, Ashworth Laboratories, Charlotte Auerbach Road, Edinburgh. EH9 3FL |

**Keywords:** Fair Isle, floristic survey, Great Britain, island diversity, Shetland

## Abstract

Fair Isle is a small isolated island located off the northern tip of Great Britain. Recognised internationally for rare migratory birds and important seabird colonies, the flora of Fair Isle has received far less attention. To rectify this, we present the first comprehensive floristic study of the island. A botanical survey was performed for each monad, and habitat information was collated following the NCC Phase 1 habitat survey method. These data were compiled to give a comprehensive checklist of 317 species, classified into 31 orders, 68 families and 191 genera according to APG IV. Of the total number of species, 254 are native to Great Britain and the remaining 63 are aliens. The list includes 10 species under threat, 7 nationally scarce and 1 nationally rare species. Our results reveal that even though Fair Isle is 200 times smaller than the full archipelago of Shetland, it holds a surprising one-third the number of species. The island is also notable for its complex mosaic of habitats, which include a range of communities that are rare or under threat elsewhere in the British Isles. We also provide recommendations for future monitoring to record changes in land-use and the effects of climate change.

## Introduction

Oceanic islands provide many spectacular examples of unlikely colonisation events and adaptation to challenging conditions. An important reason why islands are so biologically interesting is that their variation in size, shape, degree of isolation and ecology make them excellent natural laboratories that allow researchers to study ideas, generate hypotheses, and test and make predictions about dispersal and establishment (Berry, 2009; MacArthur & Wilson, 1967). Remote islands are often under-surveyed due to their isolated nature; however, the small size of many such islands makes it possible to compile full biological inventories and gain a comprehensive understanding of their biodiversity. To fully understand island biodiversity requires surveys of all groups of island biota, not just charismatic birds and mammals, with floristic surveys being particularly valuable for understanding habitat diversity. These surveys can reveal whether remote islands harbour rare species negatively affected by development on the mainland, or if the degree of isolation is effectively filtering alien species that compete with natives elsewhere (Berry, 2009; Stace & Crawley, 2015).

The United Kingdom of Great Britain and Northern Ireland (UK) has the best documented flora in the world (Plantlife, 2014), in part due to its long history of botanical interest and in part due to the restricted species number as a product of its recent glacial history. Great Britain is surrounded by over 1,000 smaller islands and islets, with over 100 islands of greatly varying size in the sub-Arctic Archipelago of Shetland (part of Vice-County 112). The flora of Shetland is well-documented with 827 species distributed across 15 habitat types. However, much of Shetland’s native flora is threatened by grazing, drainage and habitat change (P.V. Harvey, pers. comm.; W. Scott, pers. comm.; Scott & Palmer, 1987).

Fair Isle is a small, remote island half way between Orkney (42 km away) and Shetland (39 km away) (see Figure 1), though politically belongs to Shetland. The island is 5 km (3 miles) long by 2.5 km (1.5 miles) wide, with an area of 768 hectares. Its oceanic climate is greatly influenced by its small size and location, close to a major offshoot of the North Atlantic Current (NAC), and the island is one of the windiest lowland sites in the British Isles. Despite its remote nature and small size, Fair Isle is important for its biodiversity as well as for attracting visitors. It is internationally recognized as a seabird breeding site and stop-over place for migrant birds (Archer et al., 2011; Harvey & Nason, 2015). It has rugged and attractive scenery that attracts many visitors, with large areas of the island entirely unspoilt, while other parts are used for traditional crofting.

**Figure 1.**
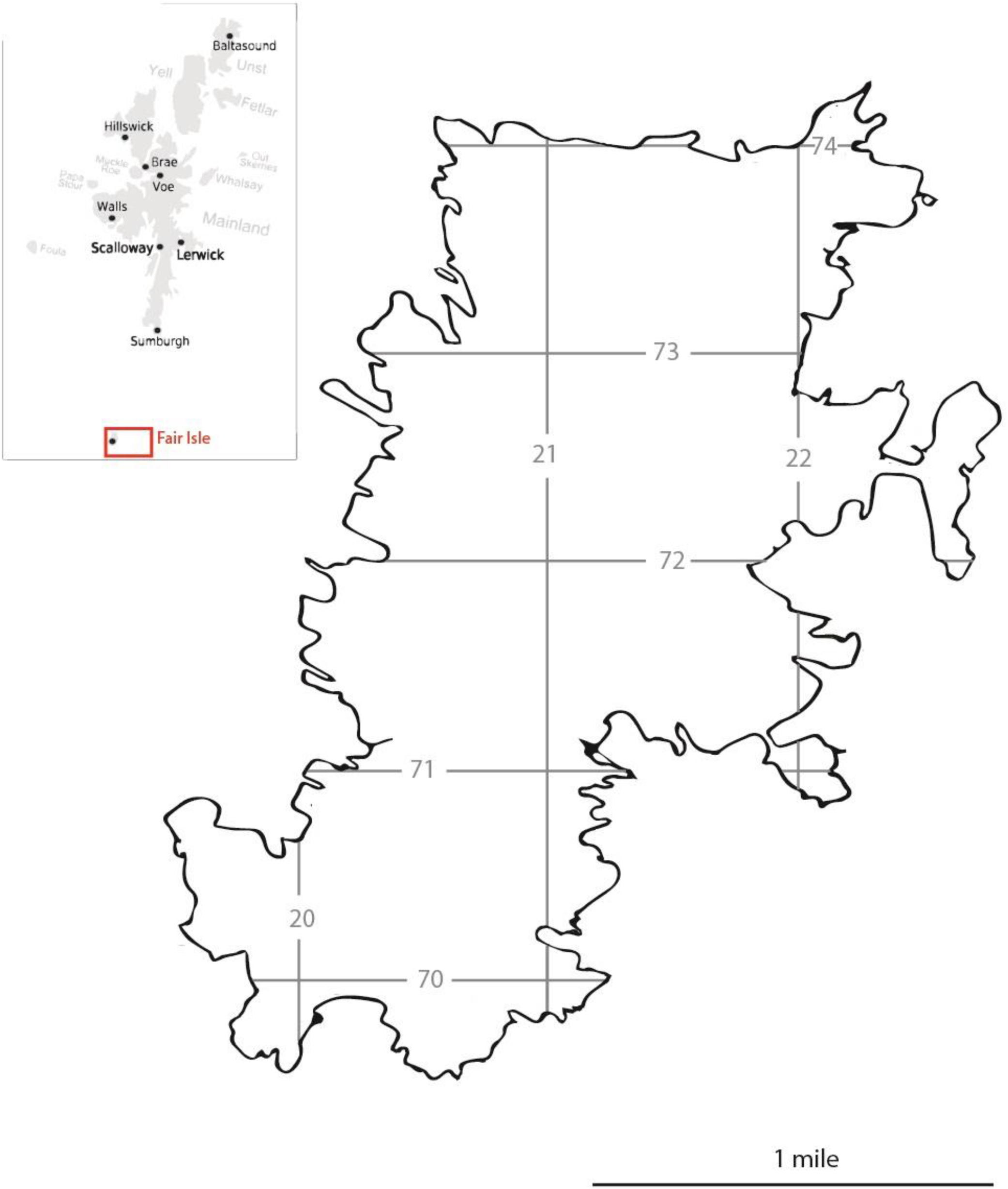
Map showing of Fair Isle, and location within Shetland.

Fair Isle has a long but intermittent history of botanical surveys. The first published floristic work was produced by James Trail in 1906, who compiled a list of 136 species made by previous observers. Pritchard (1956) contributed to our understanding of the botany of Fair Isle in a paper listing 183 plant species. This included 48 plant species not recorded by Trail, while 26 species recorded by Trail were not seen by him. Fitter (1959) reanalysed previous records and made a new island species list of 64 plant species. Currie (1960) published detailed notes on the plants of Fair Isle, reporting 12 new records and confirming many previous observations. Palmer & Scott (1963), published an account with 24 new records, but cast doubt on some species observed by Fitter (1959). The last exhaustive published study of the botany of Fair Isle was by Walter Scott (1972), whose account included some 238 taxa. This included some doubtful species in need of verification, and a list of 15 widespread species found commonly across the rest of Shetland but absent from Fair Isle. An island-wide habitat survey in 1991-92, recorded 218 taxa, including eight species known to be planted and one hybrid, with three species, *Spergularia marina, Drosera rotundifolia* and *Mertensia maritima*, additional to those listed in Scott (1972) (Riddiford, unpublished manuscript, 1992). A further study, in 2002, used the National Vegetation Classification (NVC) methodology to describe habitats (O’Hanrahan, 2003) but this was limited to the Site of Special Scientific Interest (SSSI) in the northern part of the isle. While relatively comprehensive in their coverage, none of these historical botanical studies give detailed accounts of the distribution of species across the island, nor directly relate this to the diversity of habitats.

The current study looks to update and consolidate previous work, while filling the knowledge gap in habitat diversity. The first major aim is to assess the flora of Fair Isle through a qualitative floristic survey. This will allow the generation of a comprehensive checklist of the Flora of Fair Isle, updated with the classification of the Angiosperm Phylogeny Group IV (APG IV) (Byng *et al*., 2016). We also consider how many species are native, introduced, under threat, nationally rare and scarce, and compare these observations to other islands. Secondly, we endeavour to identify the various habitats present on the island using the Nature Conservancy Council (NCC) Phase 1 habitat categories, and link these habitats to observations of species diversity. Overall, these results will give key insights into the floristic diversity of an important and potentially biologically diverse remote island.

## Methods

### Field survey

The project involved a comprehensive floristic survey of the island in monads, recording all angiosperms, gymnosperms, lycopodiophyta and pteridophyta. The survey methodology of recording in one km grid square was the same as that used for plant recording by the Botanical Society of Britain and Ireland (BSBI), so that these records could contribute to the Atlas of the British and Irish Flora, which is currently in the process of being revised. Monad boundaries followed the Ordnance Survey map for Shetland, as shown in Figure 1. All 19 monads were included in the survey (see Supplementary Text S1 for habitat description and histories for each of the 19 monads surveyed). Informal botanical surveys were made from 1981 to present, then a comprehensive island-wide survey for each monad was made in June 2016. Most identifications followed Poland (2009), Rose (1989) for grasses, rushes and sedges (Poaceae, Juncaceae and Cyperaceae respectively) and Streeter (2009). These data were added to historical records deposited in the BSBI Distribution Database BSBI (2015), and the published accounts of Currie (1960), Fitter (1959), Pritchard (1957), Scott (1972) and Trail (1906).

### Habitat survey

Habitat classification of each monad was made in 1991-1992 following the guidelines for NCC (1990) Phase 1 habitat surveys. Fieldwork was conducted in June to August in both years to coincide with the flowering season. Mapping was done directly in the field and an assessment made simultaneously of the dominant and frequent contributors to the plant community in each habitat. Where two habitats were intermingled, they were treated as habitat mosaics, a category additional to the NCC protocol; this particularly applied to some areas of grassland and heath, labelled grassland/heath mosaic. The only other deviation from the protocol involved the incorporation of prominent bryophyte and lichen communities into the habitat assessment. Each part of the island was visited at least once, with repeat visits made to clarify any uncertainties and to confirm habitat boundaries from original draft maps. Once the draft map was completed, the boundaries were transferred to 1:2500 Ordnance Survey maps and the area for each habitat calculated by overlay of squares of known size ratio to achieve an approximate figure. Additional habitat observations were made in June 2016 to check for habitat changes since the 1991/92 survey.

### Interpretation of the data

A database of species presence/absence was made, in a file modified from the BSBI format to enter records from a Biological Record Centre (BRC) Recording Card. These data used the taxonomy of APG IV (Byng *et al*., 2016) and The Plant List (2013), Tropicos (2016) for synonymy and IPNI (2012) for authority. We used Stace & Crawley (2015) to infer the native/alien status, with alien taxa further divided into archaeophytes (AR; those taxa already present in the UK prior to the year 1500) and neophytes (AN; an alien plant that has arrived in the UK since the year 1500). We used the BSBI website to infer national rarity and scarcity, where rare plants are those recorded in 1-15 ten km squares and scarce plants those recorded in 16-100 ten km squares. We used Cheffings *et al.* (2005) to infer the national red list status. In addition to these classifications, we assigned each species a ‘Fair Isle status’ to highlight abundance on the island. Categories were: Native Extant, Native Extinct, Native Recent Colonist, Natural Colonist Established, Natural Colonist Failed/Short-lived, Casual Established, Casual Temporarily Established, Casual Extinct, Casual Failed Short-lived, Planted Established, Planted Temporarily Established, Planted Failed/Short Lived, Escapes, Unverified and Doubtful (see Supplementary Table S2 for categories of Fair Isle status).

## Results

### Field survey

The full species checklist included a total of 317 vascular plant species recorded across the island (Table 1). These species include representatives from five plant divisions (pteridophyte, 11 species; lycopodiophyta, 2 species; gymnosperms, 4 species; angiosperm-monocots, 86 species; angiosperm-eudicots, 214 species), and a total of 31 orders, 68 families and 191 genera. The families of angiosperms with the largest number of species were Asteraceae and Poaceae, followed by Cyperaceae, Brassicaceae and Caryophyllaceae (Table 2). These five families comprise 41% of all observed species.

**Table 1.**
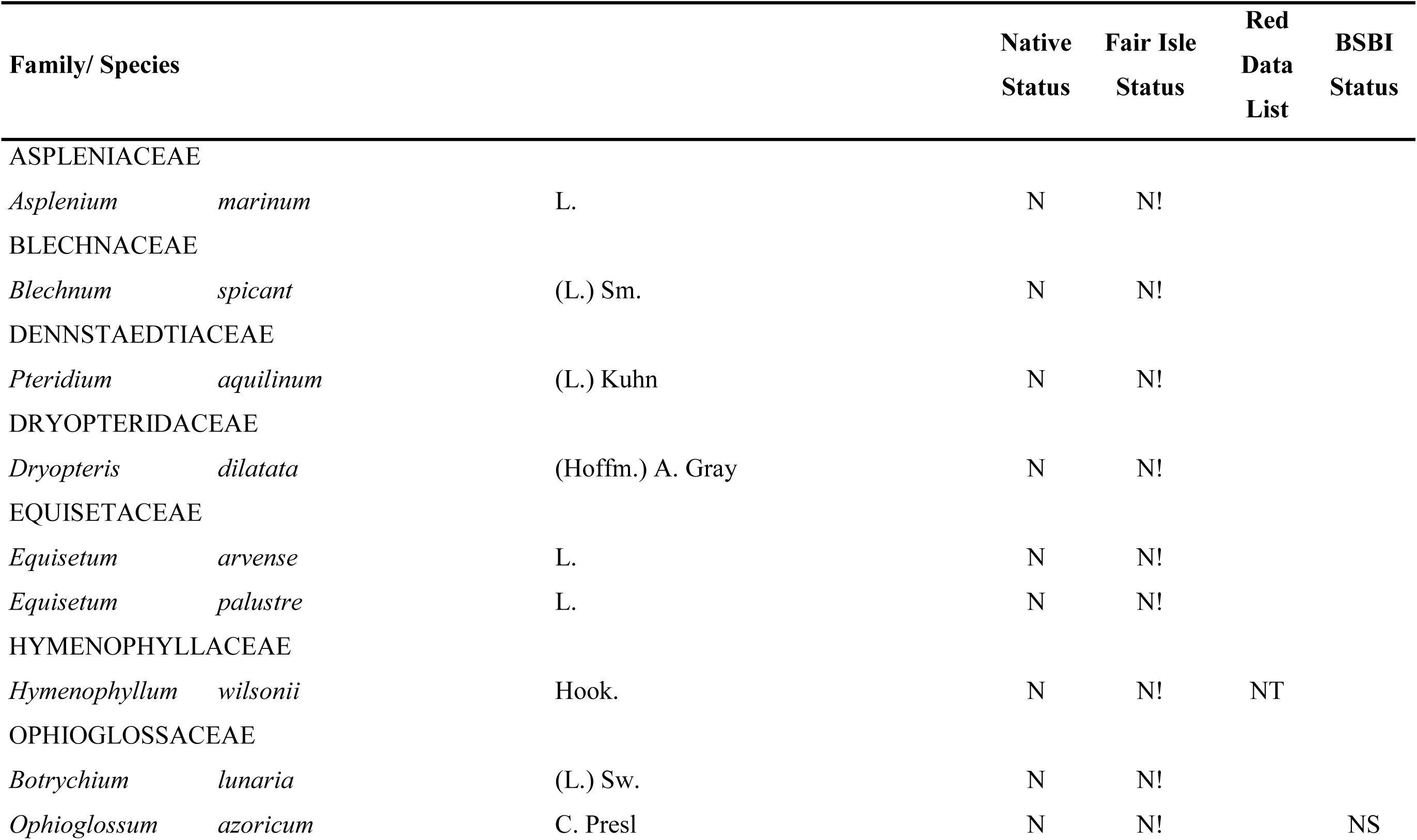
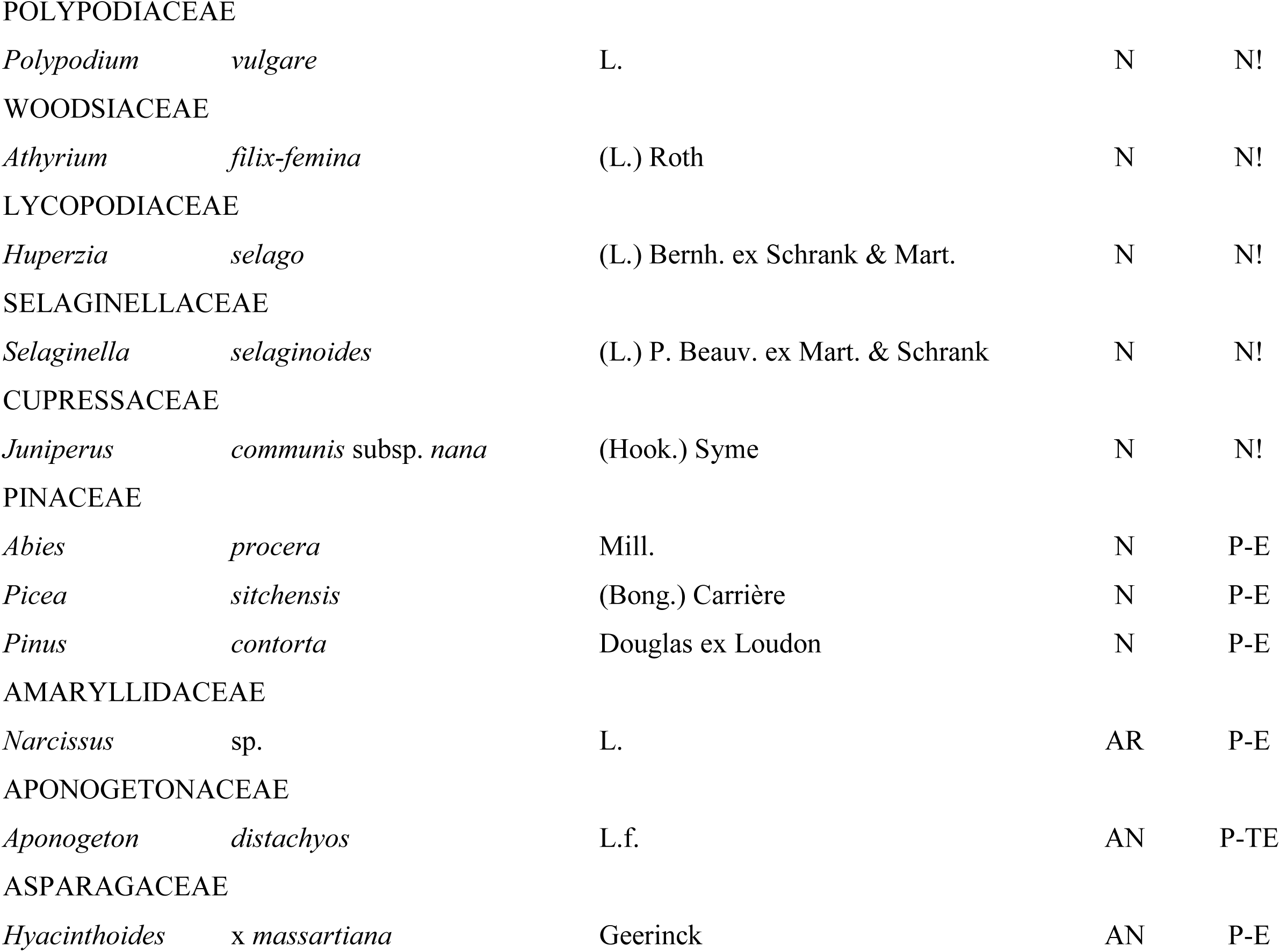
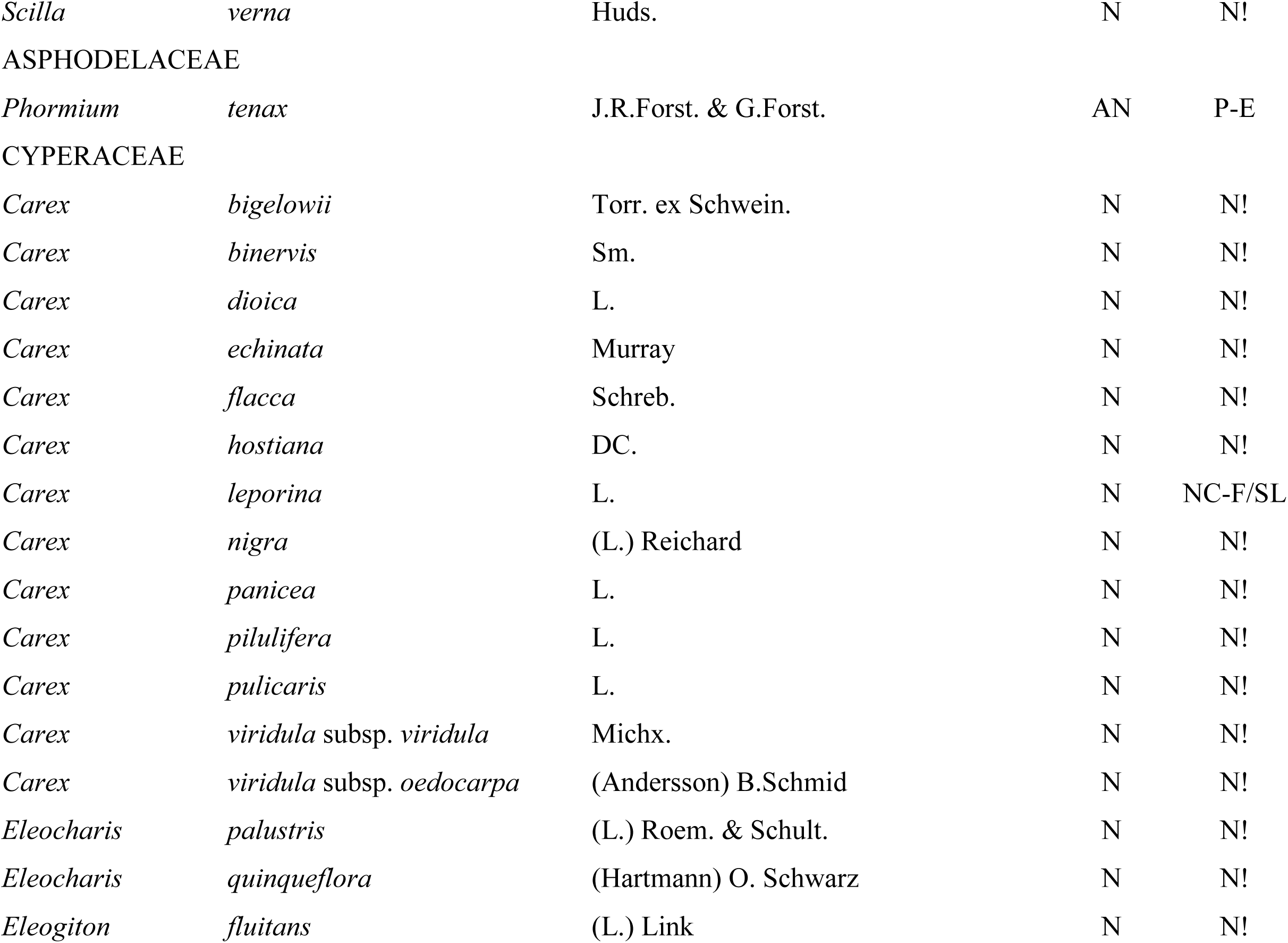
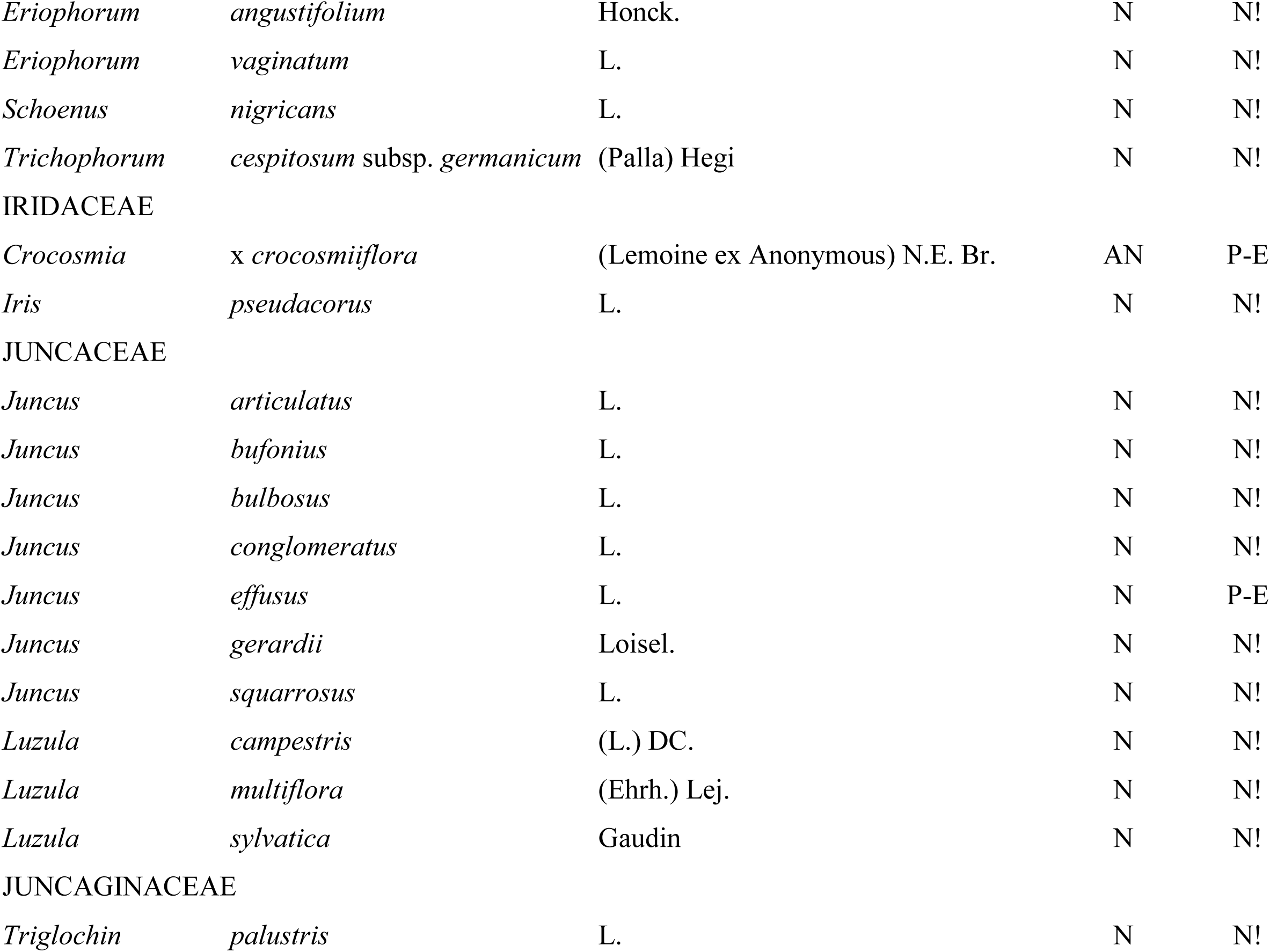
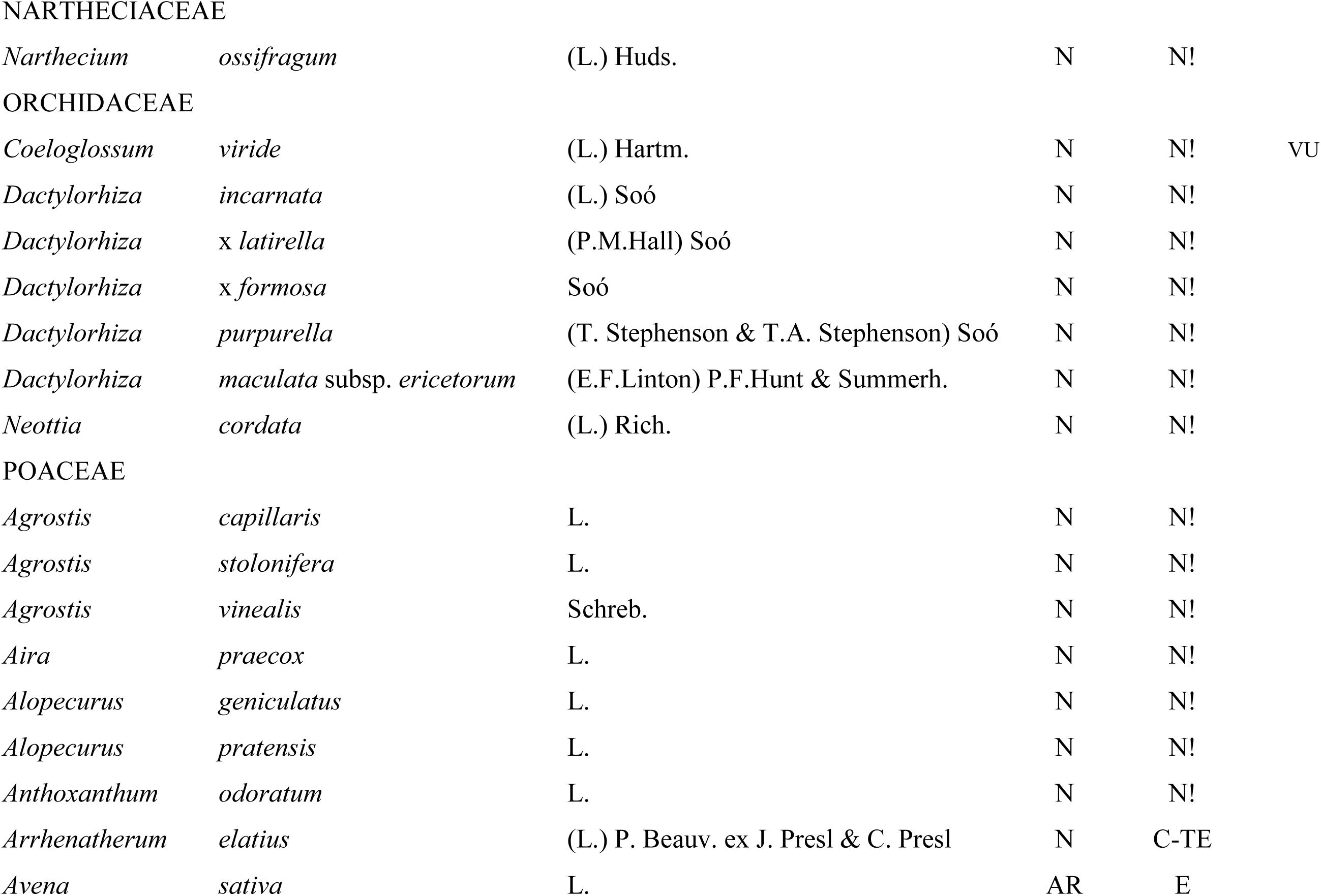
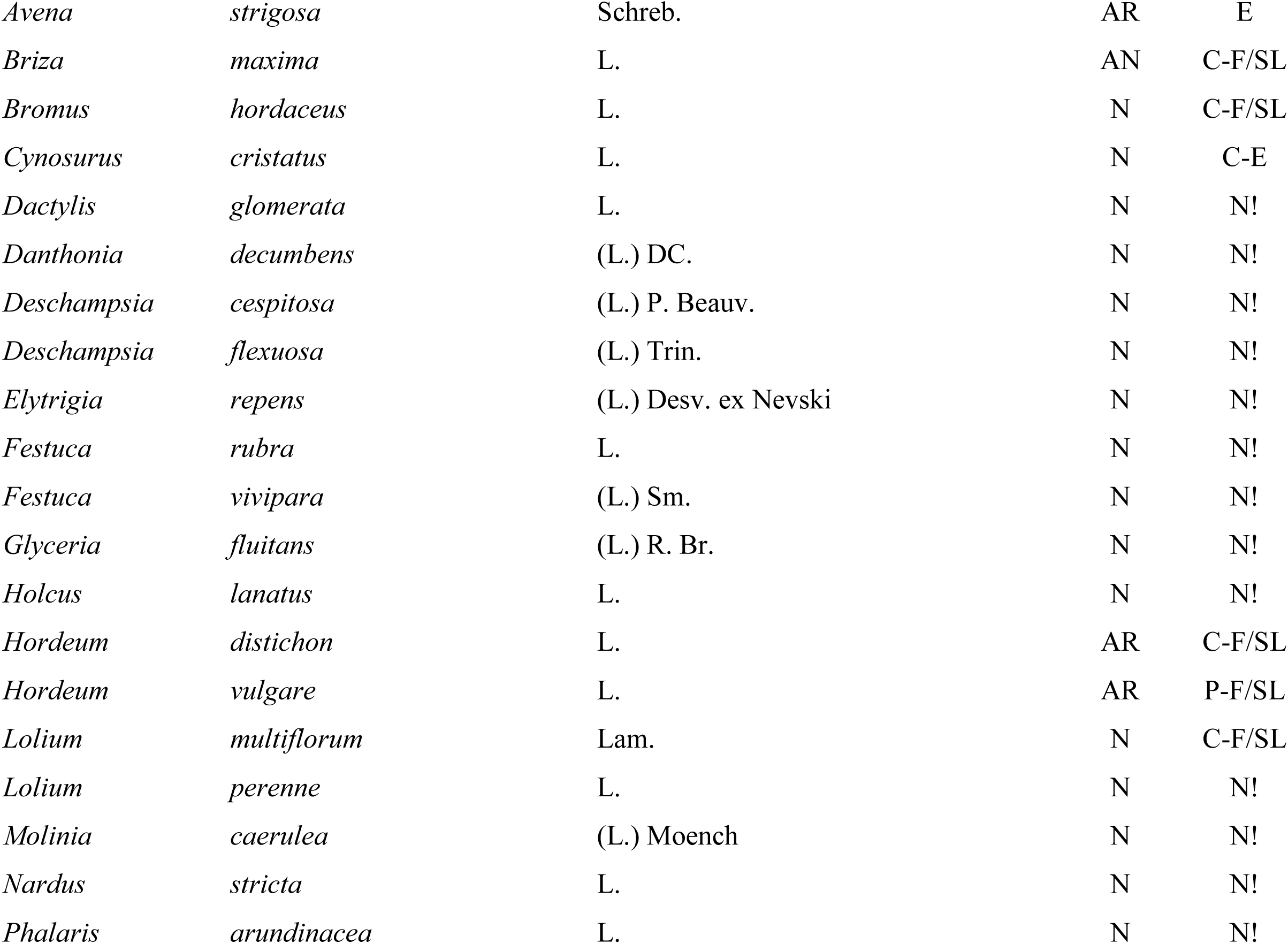
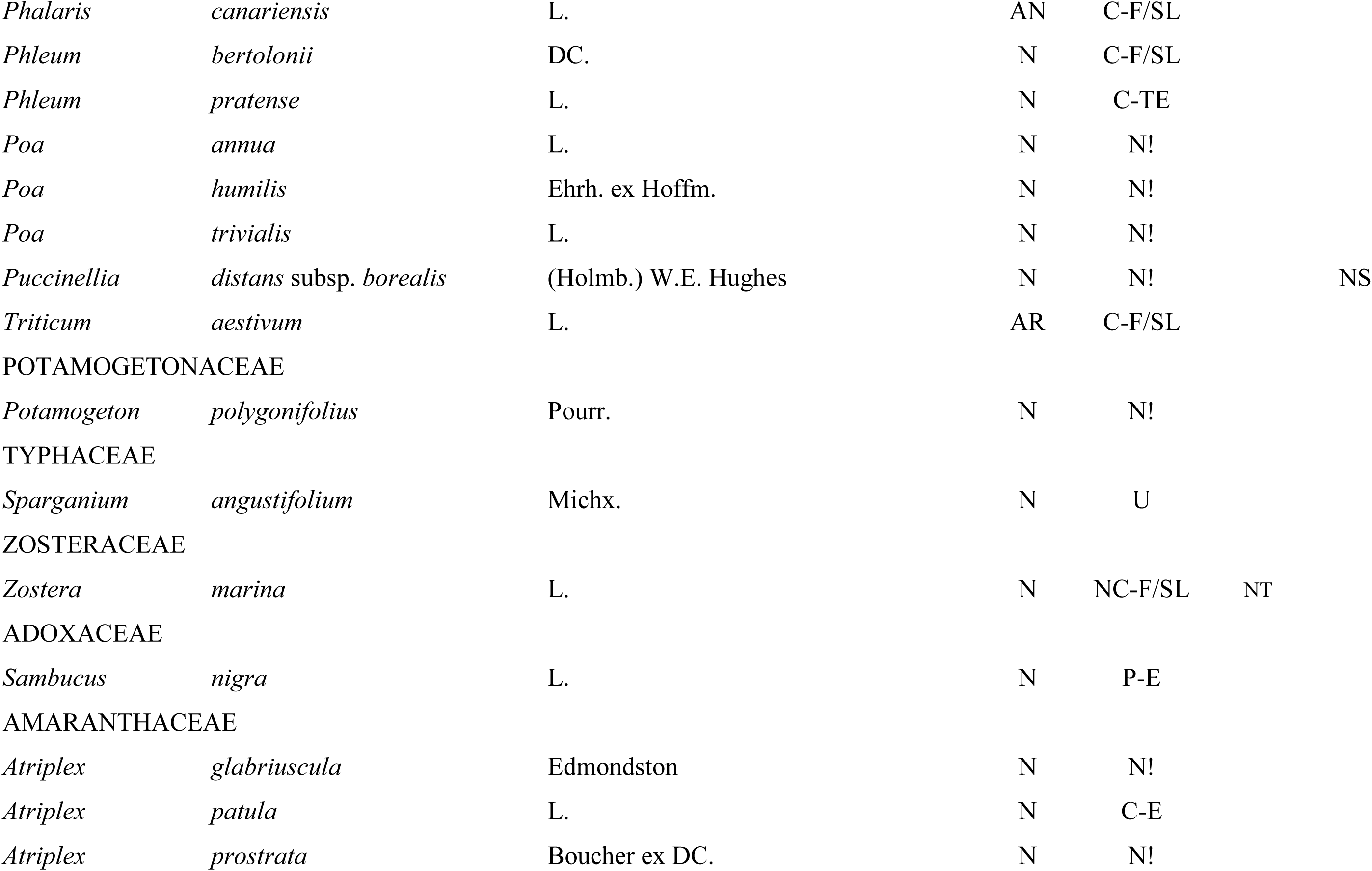
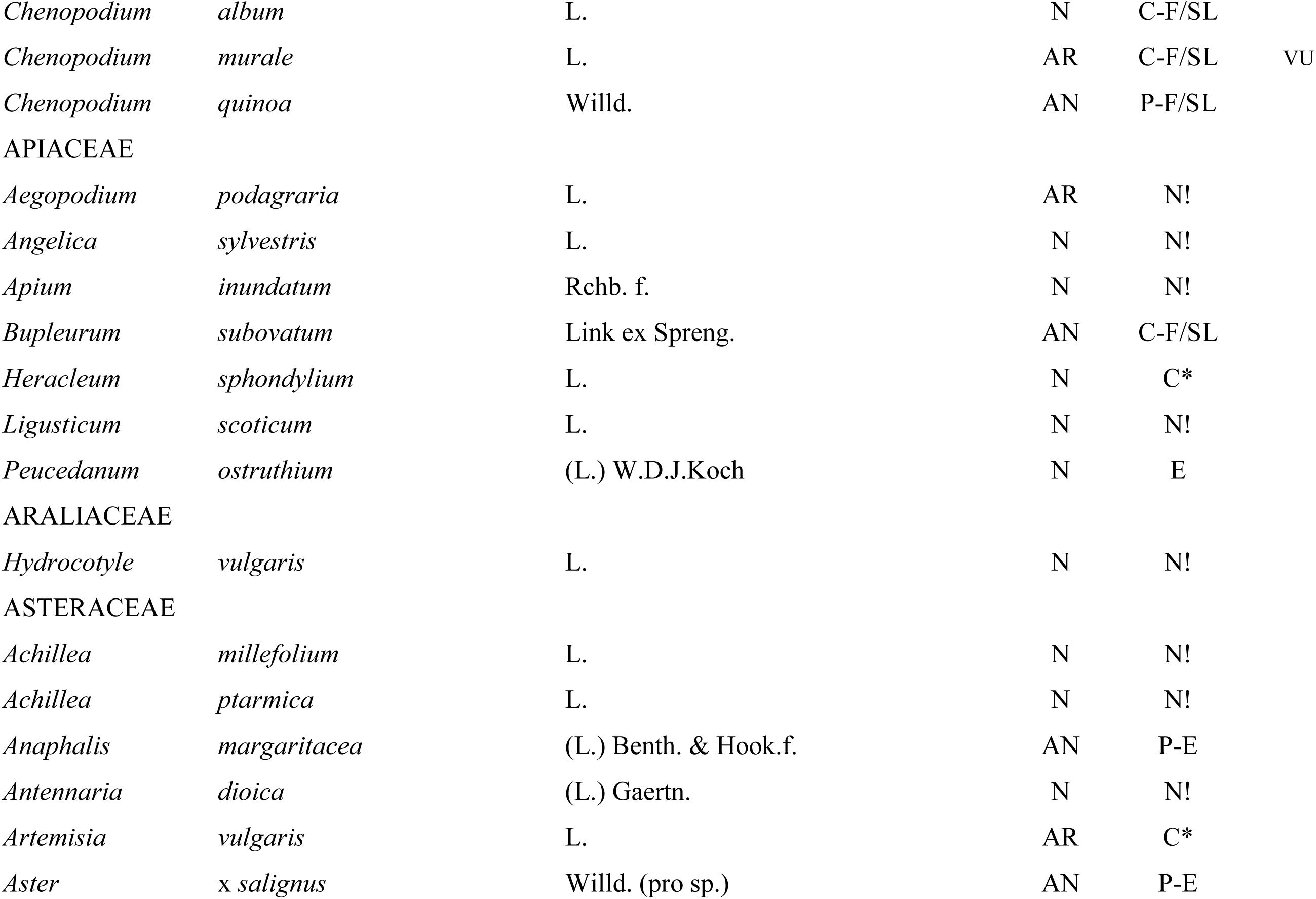
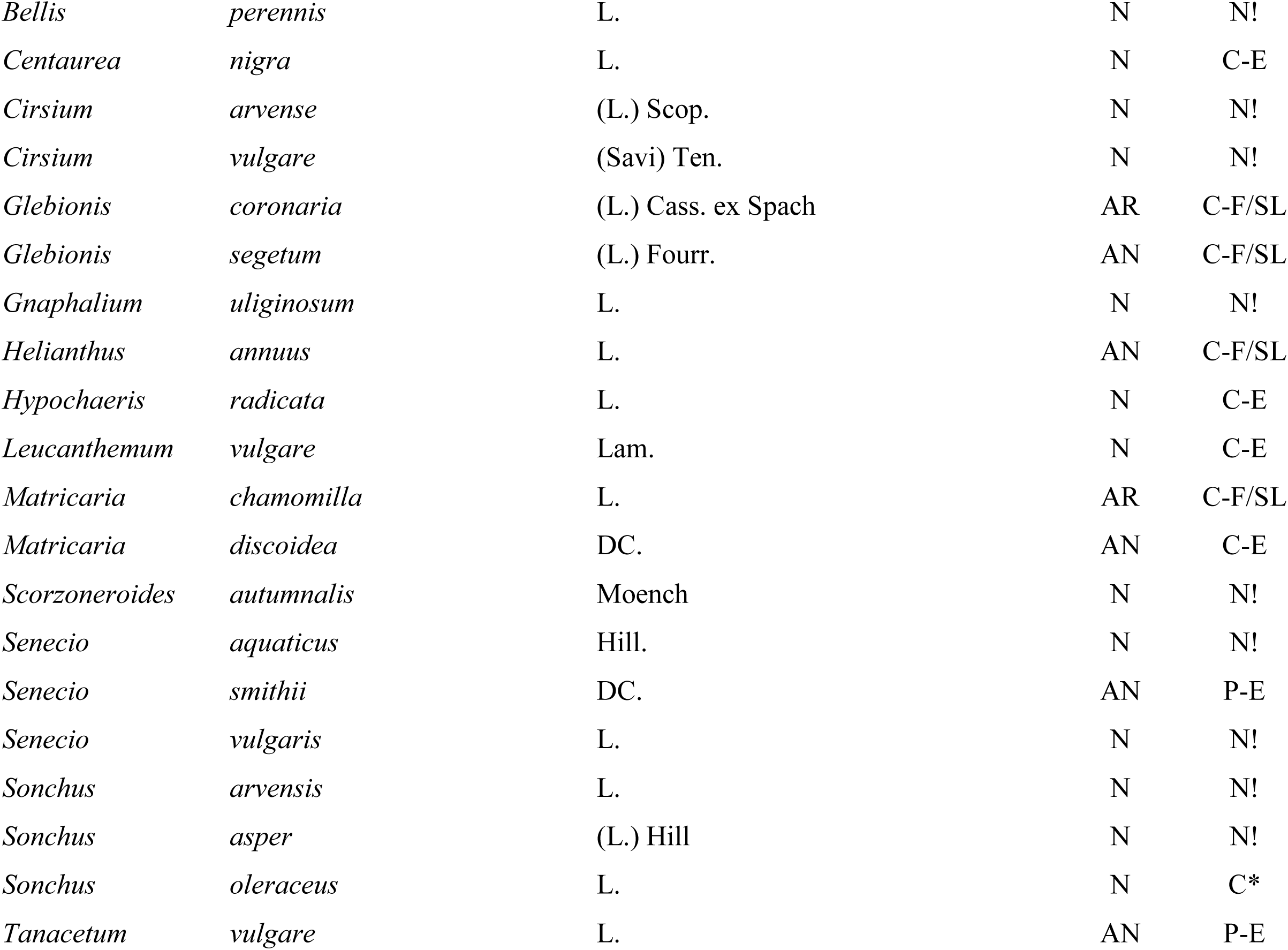
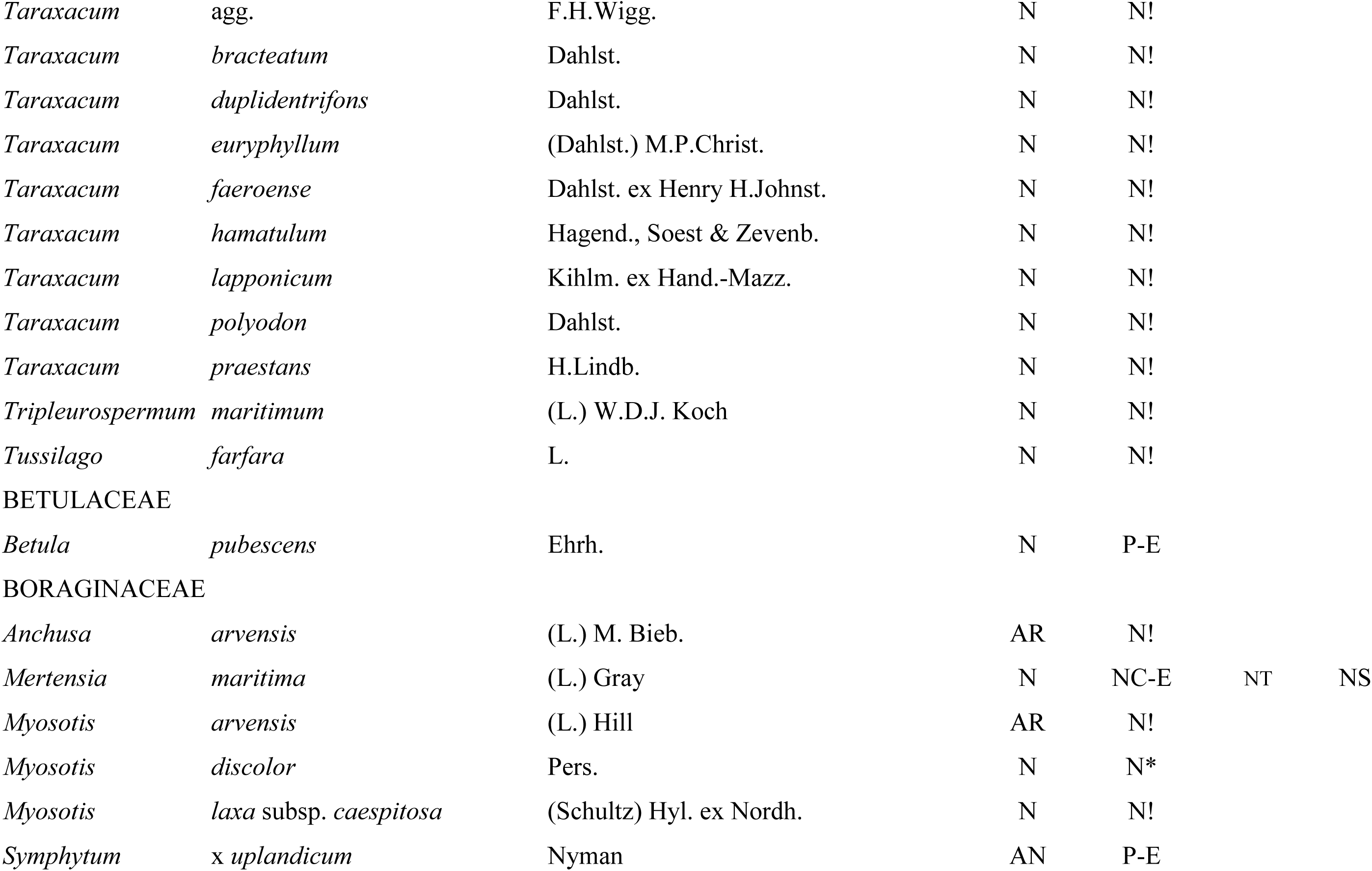
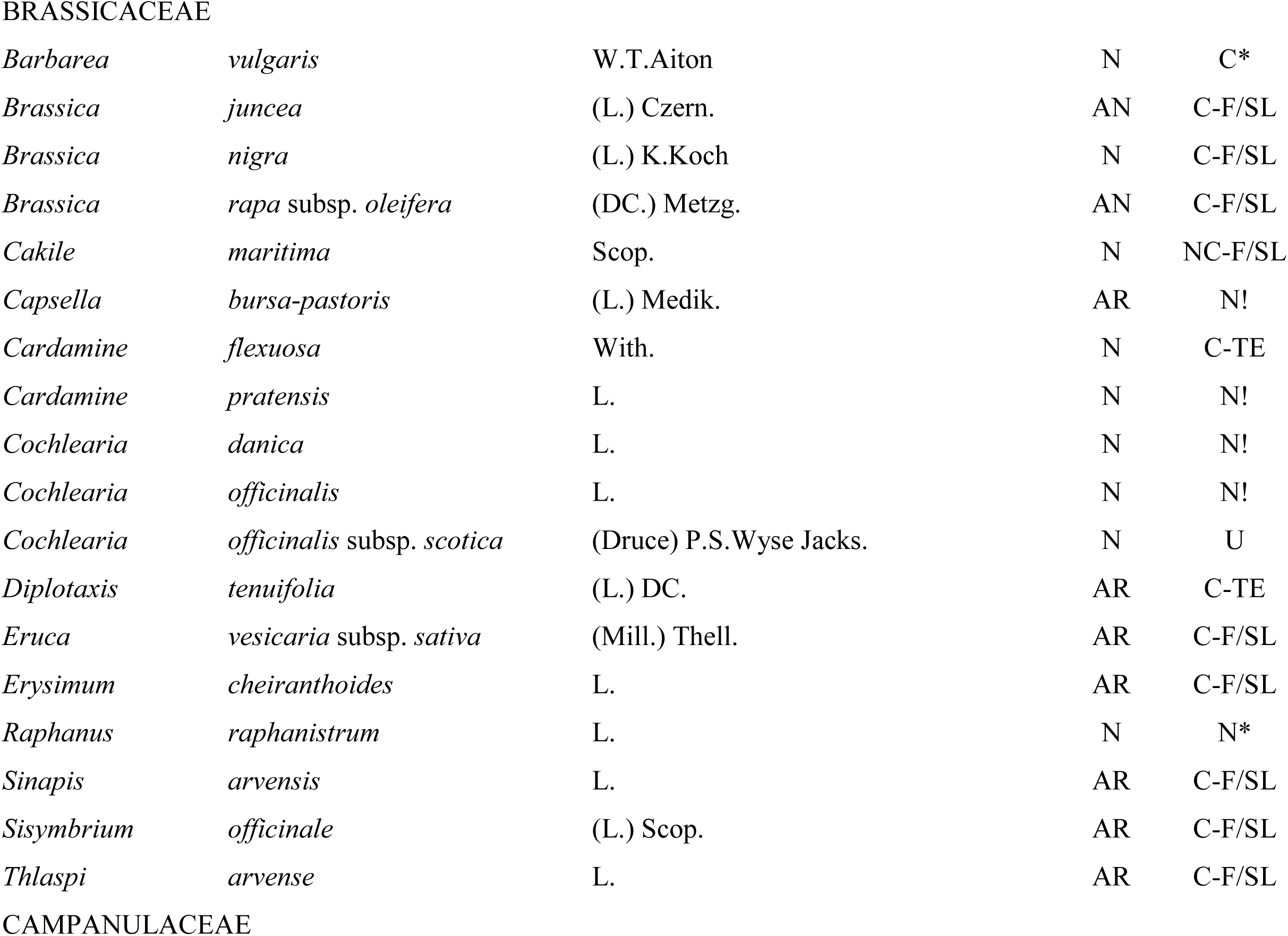
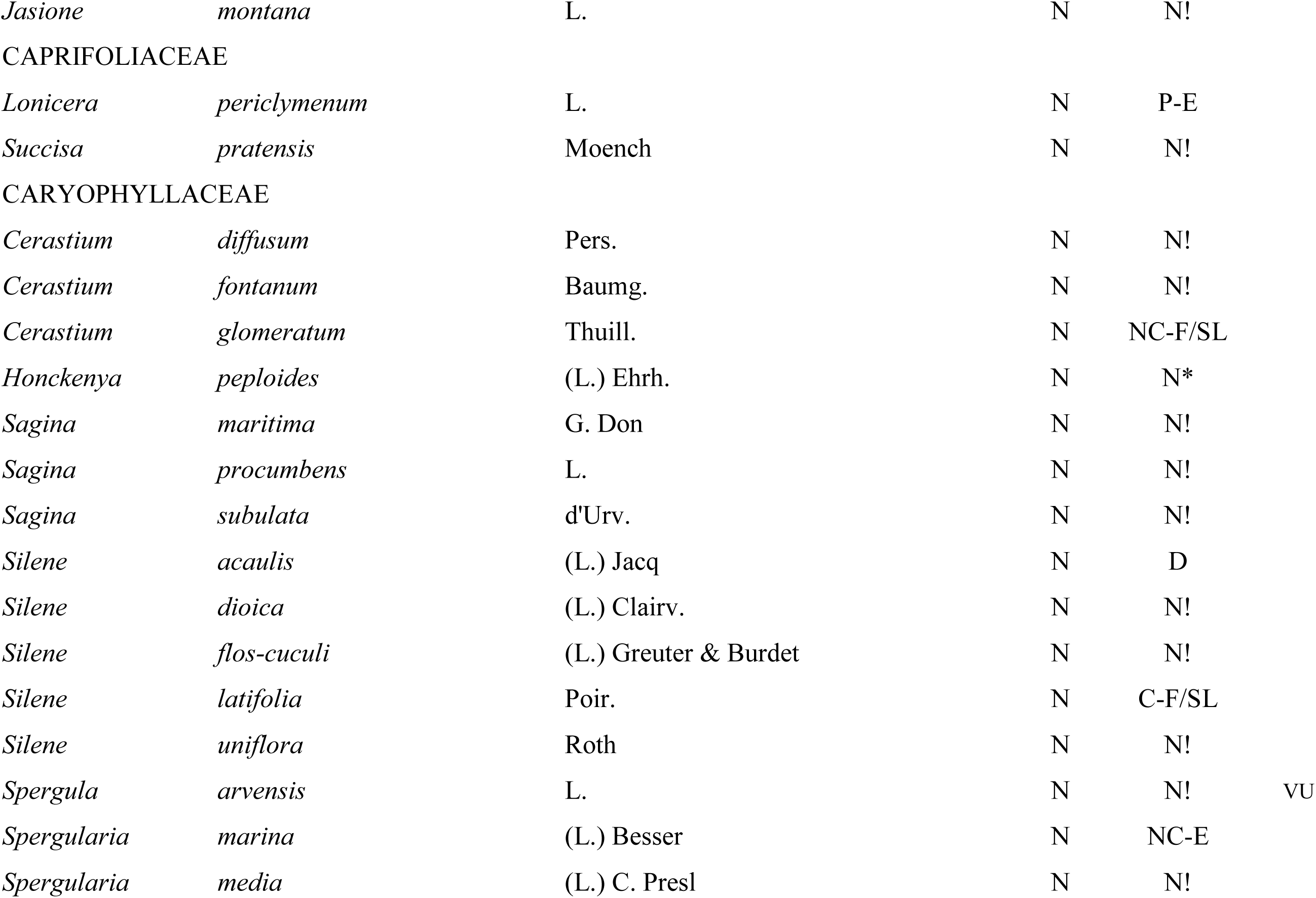
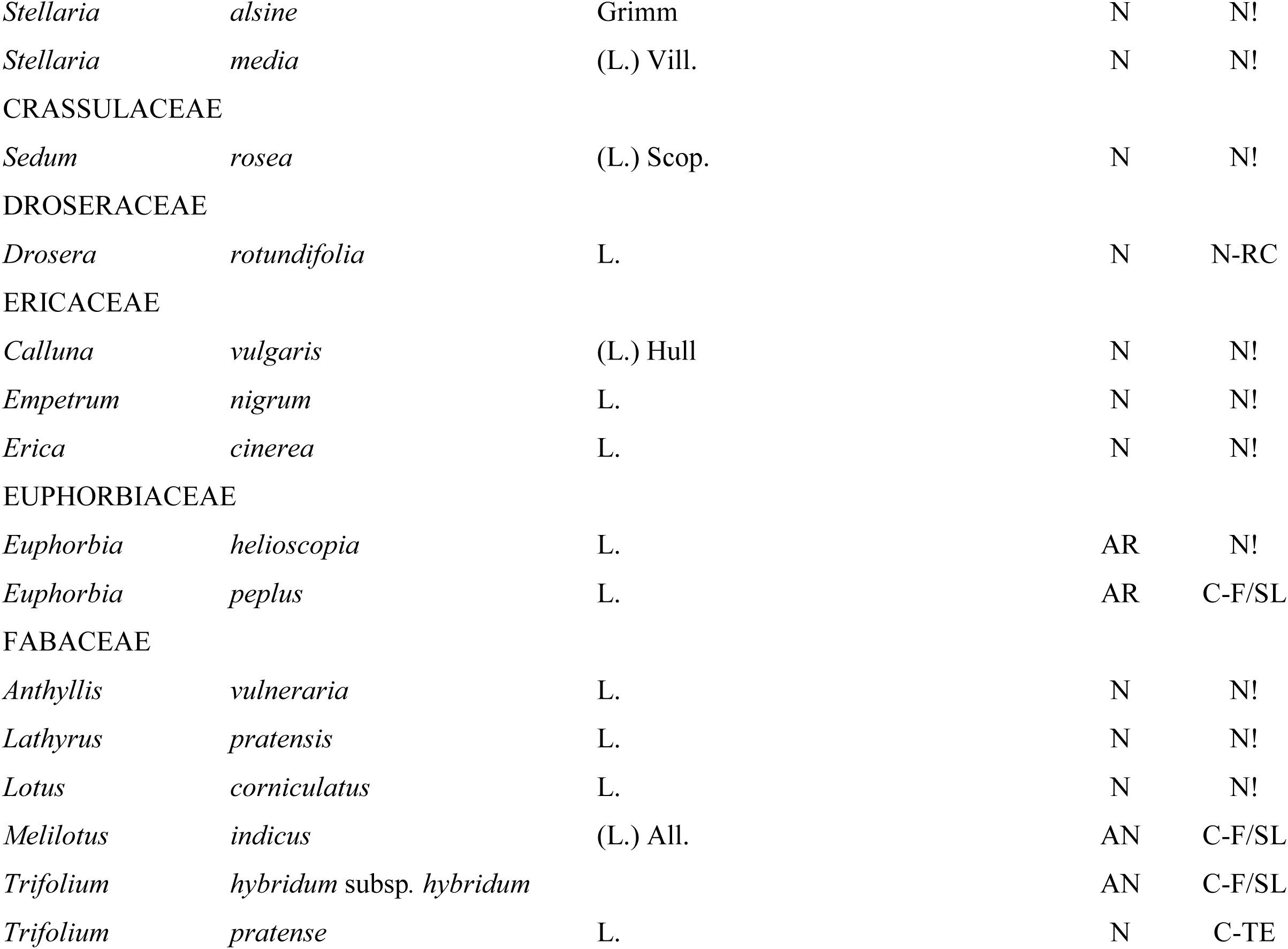
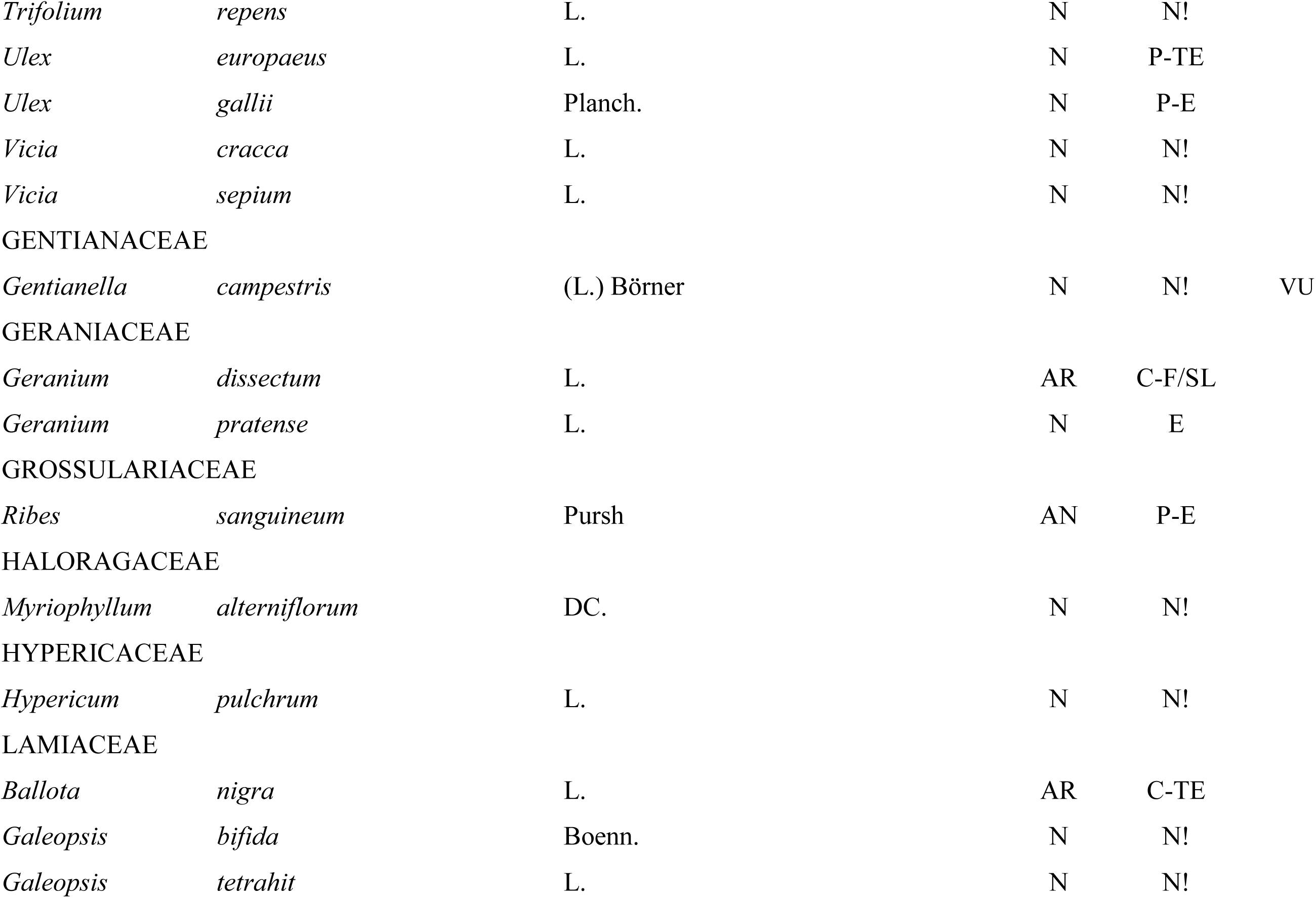
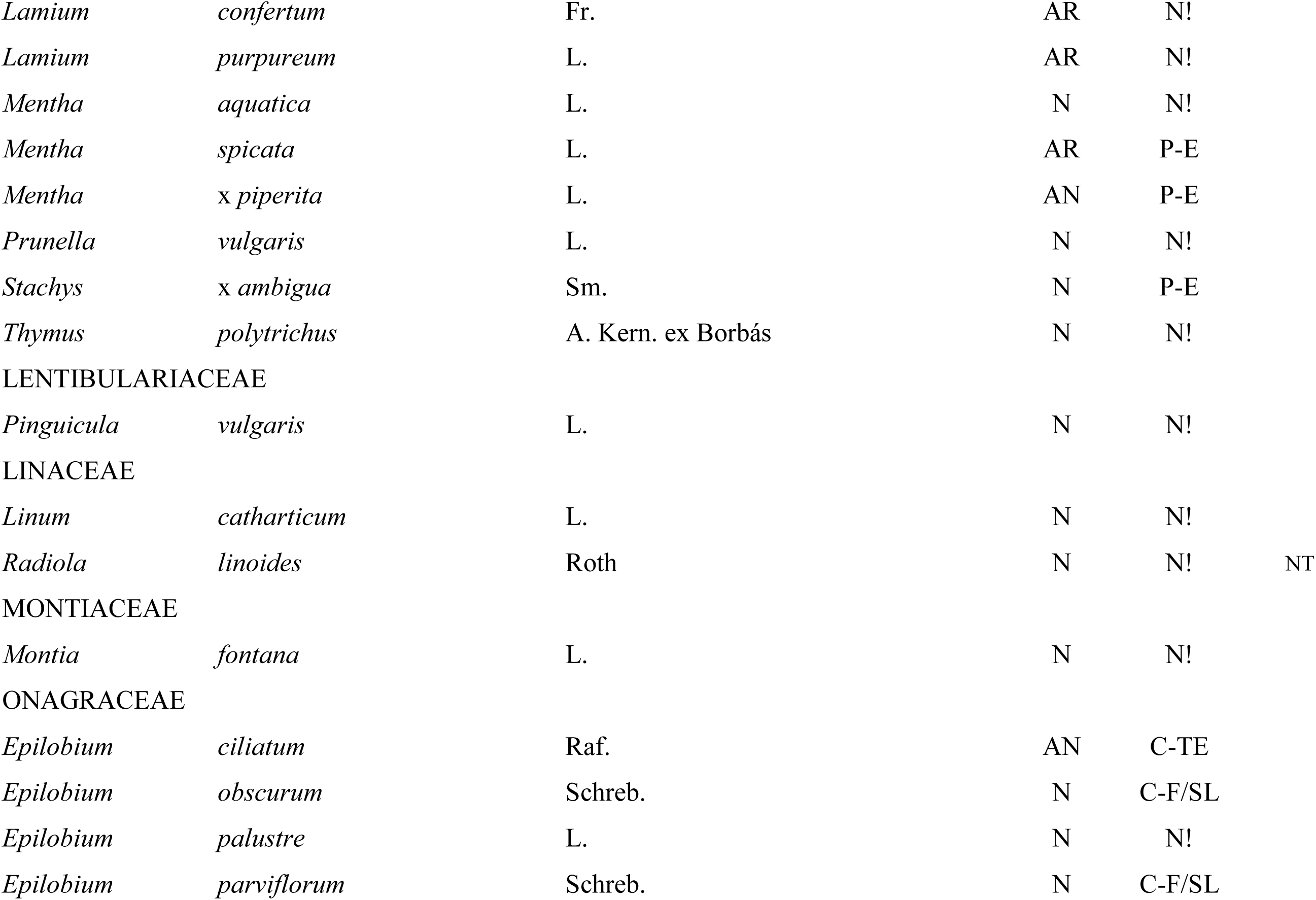
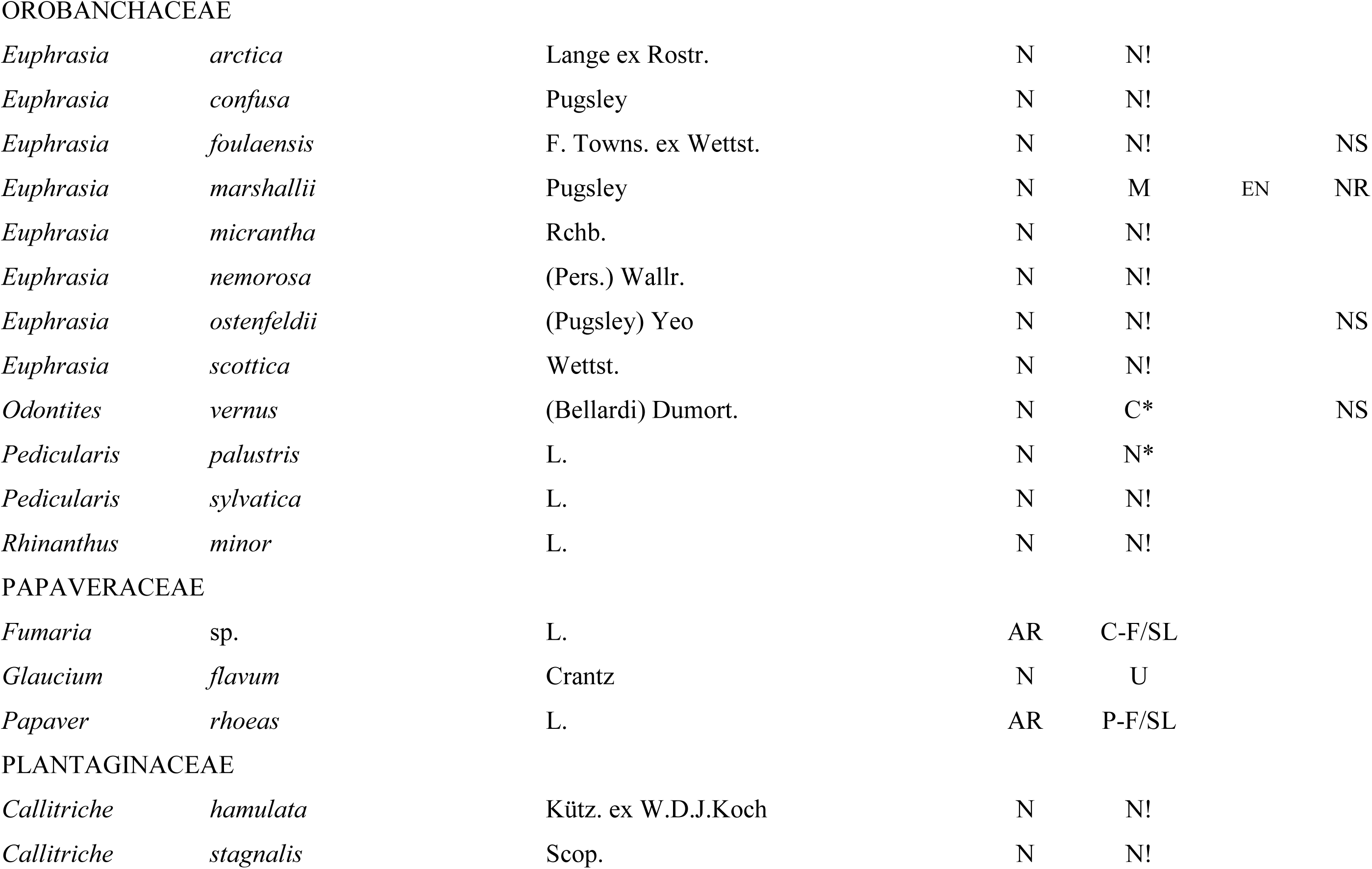
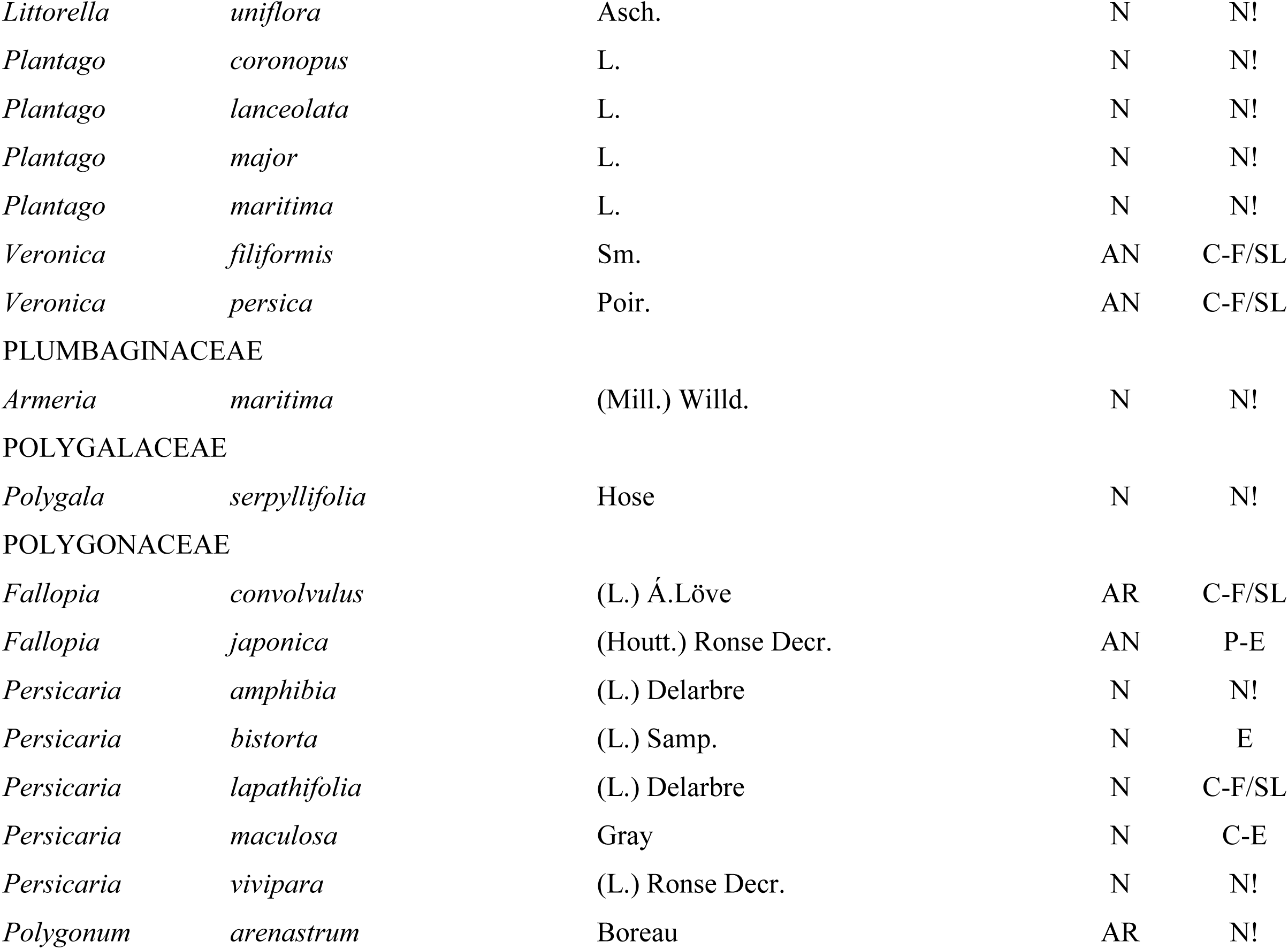
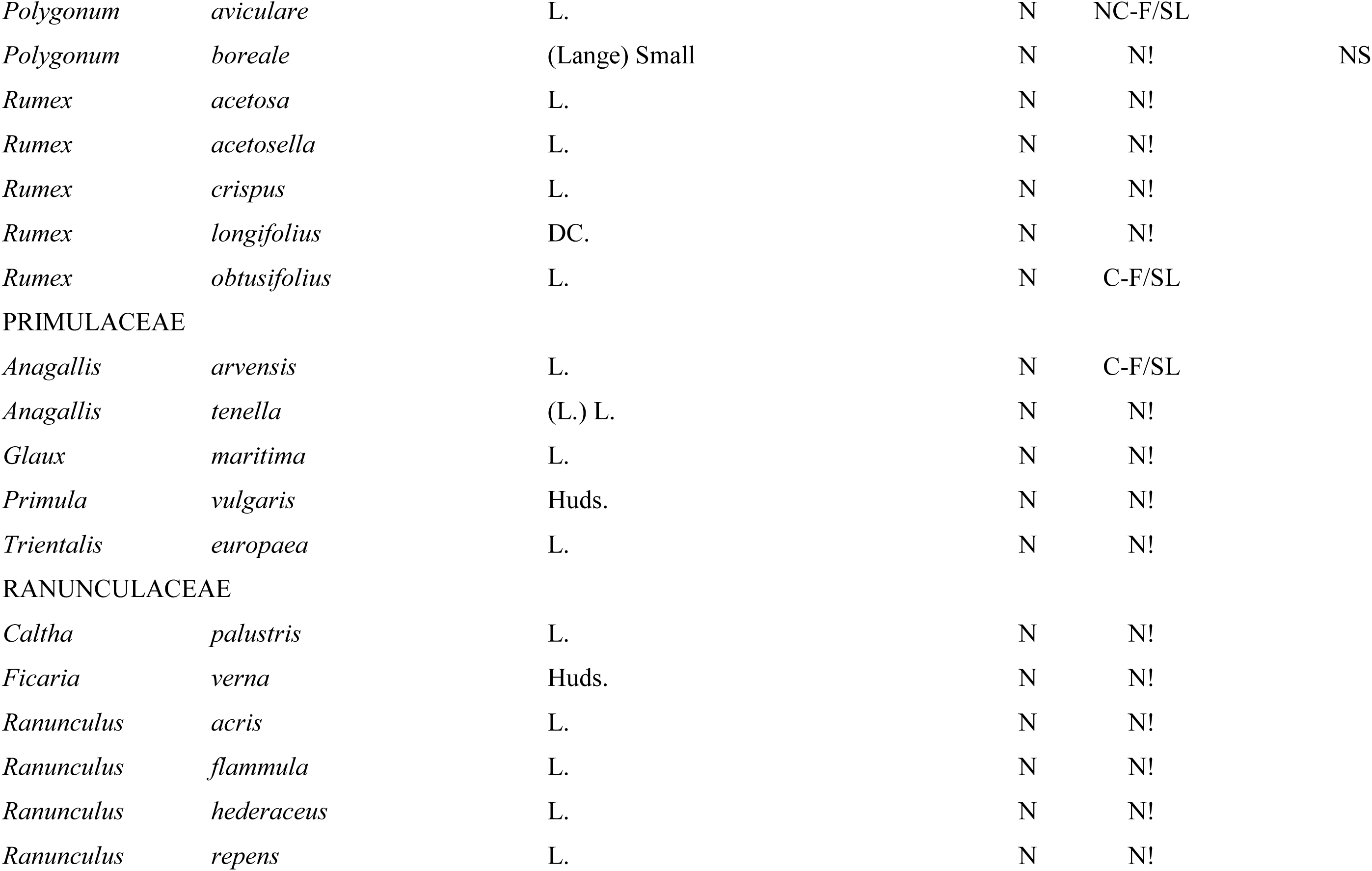
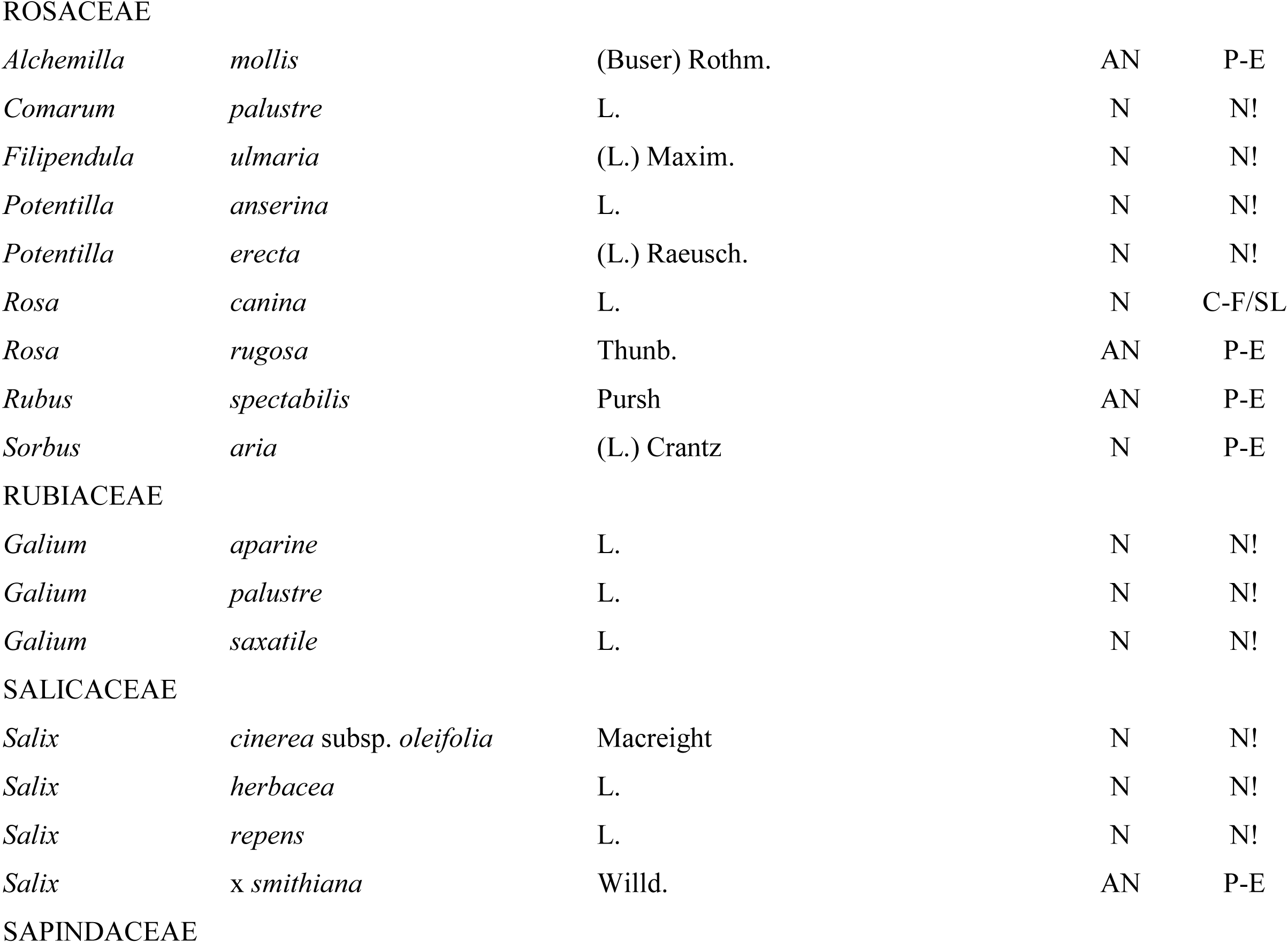
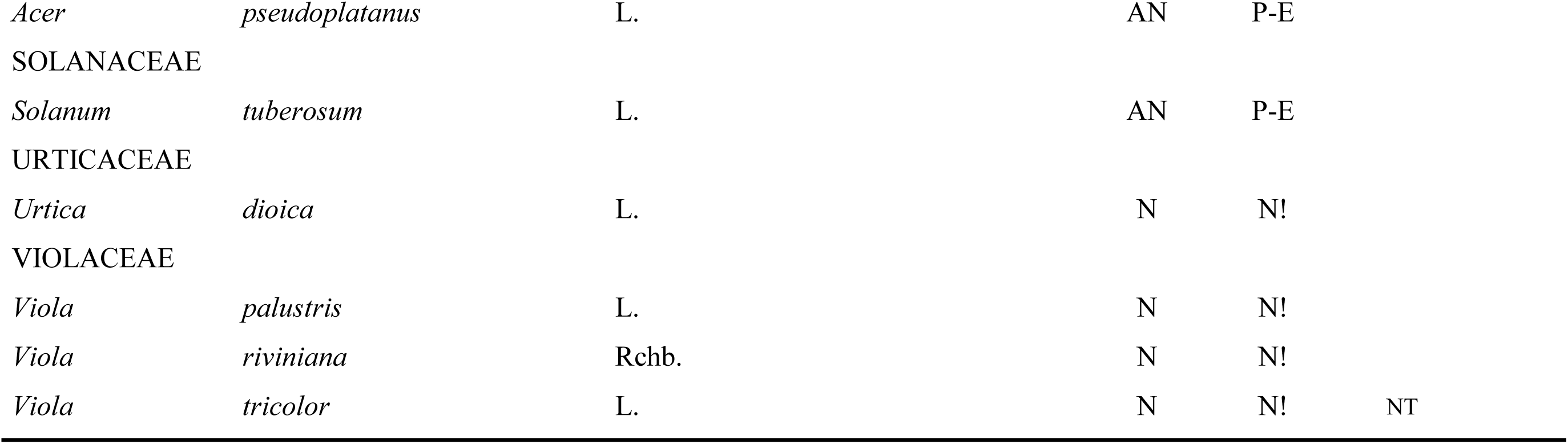
Comprehensive checklist of the Flora of Fair Isle, arranged by family following APG IV (Byng *et al*., 2016). Fair Isle Status is described in Supplementary Table S2.

**Table 2.** Breakdown of most well represented plant families on Fair Isle.

| **Ranking** | **Family** | **N° of genera** | **N° of species** | **% of all family** |
| --- | --- | --- | --- | --- |
| 1 | Asteraceae | 21 | 37 | 12 |
| 2 | Poaceae | 25 | 37 | 12 |
| 3 | Cyperaceae | 6 | 20 | 6 |
| 4 | Brassicaceae | 13 | 18 | 6 |
| 5 | Caryophyllaceae | 7 | 17 | 5 |
| Total |  | 72 | 129 | 41% |

Additions to the flora of Fair Isle made by Scott & Palmer (1987), either from new records generated in the course of this survey, or from recent literature, are reported in Table 3. *Phleum bertolonii* (as *P. pratense* subsp. *bertolonii*) was included in Scott & Palmer (1987), but that record was considered likely to be an error and thus this record is a first for Fair Isle. *Chenopodium quinoa* does not appear in Scott & Palmer (1987); it is best considered a crop rather than a casual, a category not included in Scott & Palmer. *Brassica juncea, Brassica rapa oleifolia, Erysimum cheiranthoides* and *Helianthus annuus* were not in Scott & Palmer (1987) but were added in Scott (2011). *Aster x salignus*, was originally determined as *Aster foliaceus* (Scott 1972), but re-determined by NJR as *Aster x salignus* in 2013. Finally, the seagrass *Zostera marina*, which is not known to grow on Fair Isle, has been recorded on the strand line by Nora McMillan (McMillan, 1986) and verified by Walter Scott, and thus is best treated as drift material, perhaps from the nearest colonies in Orkney (Scott, 2011).

**Table 3.**
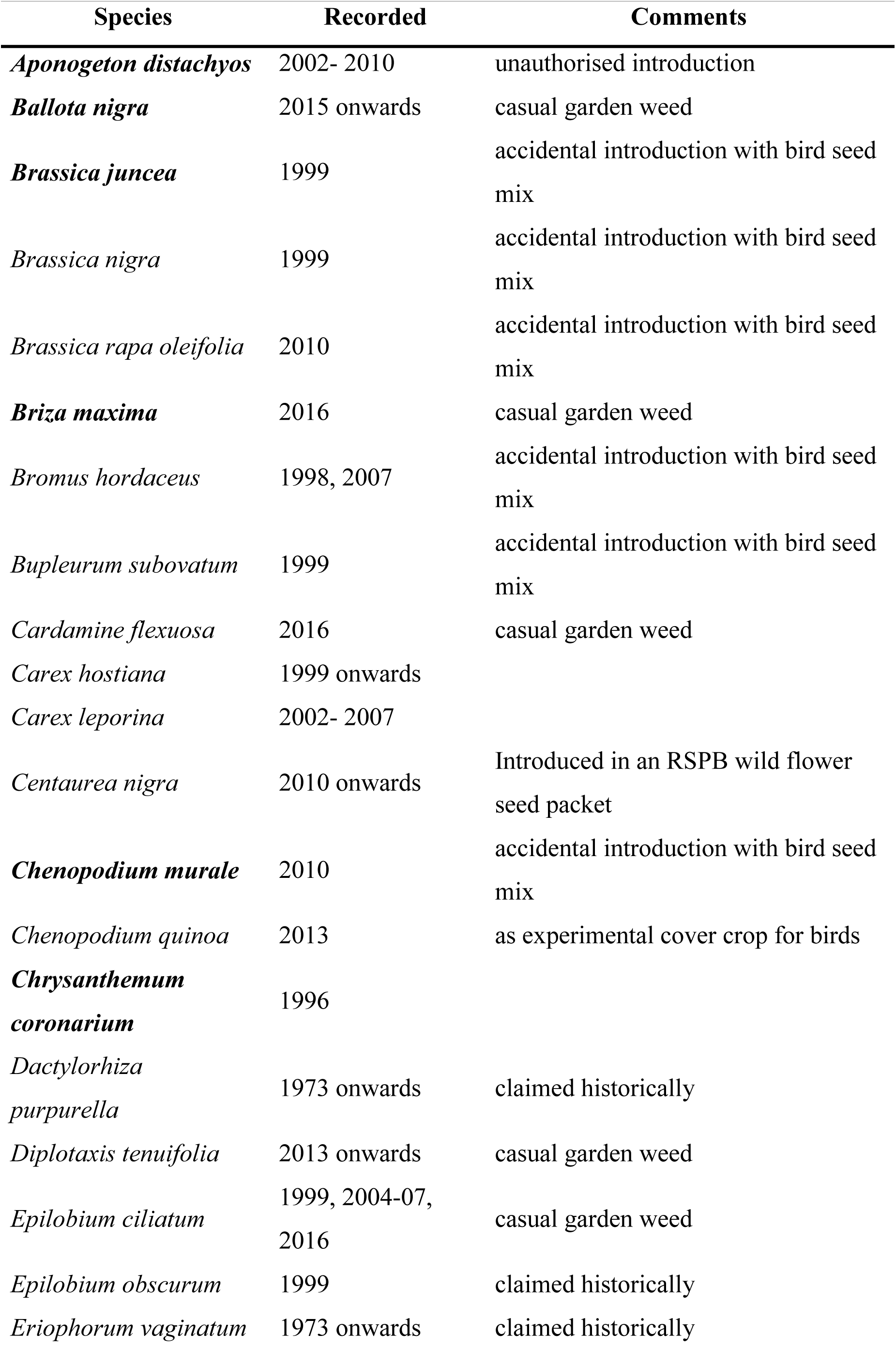
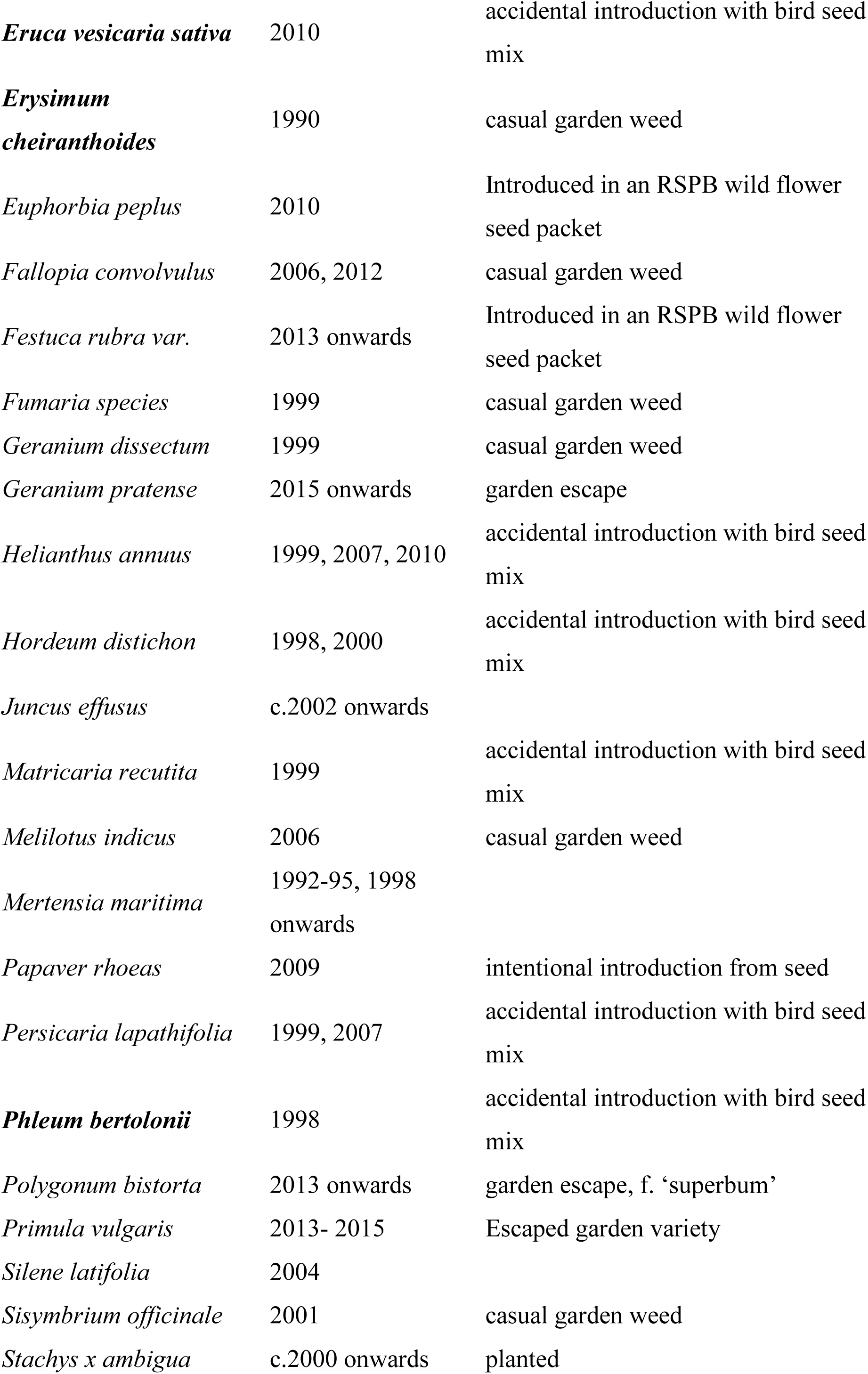
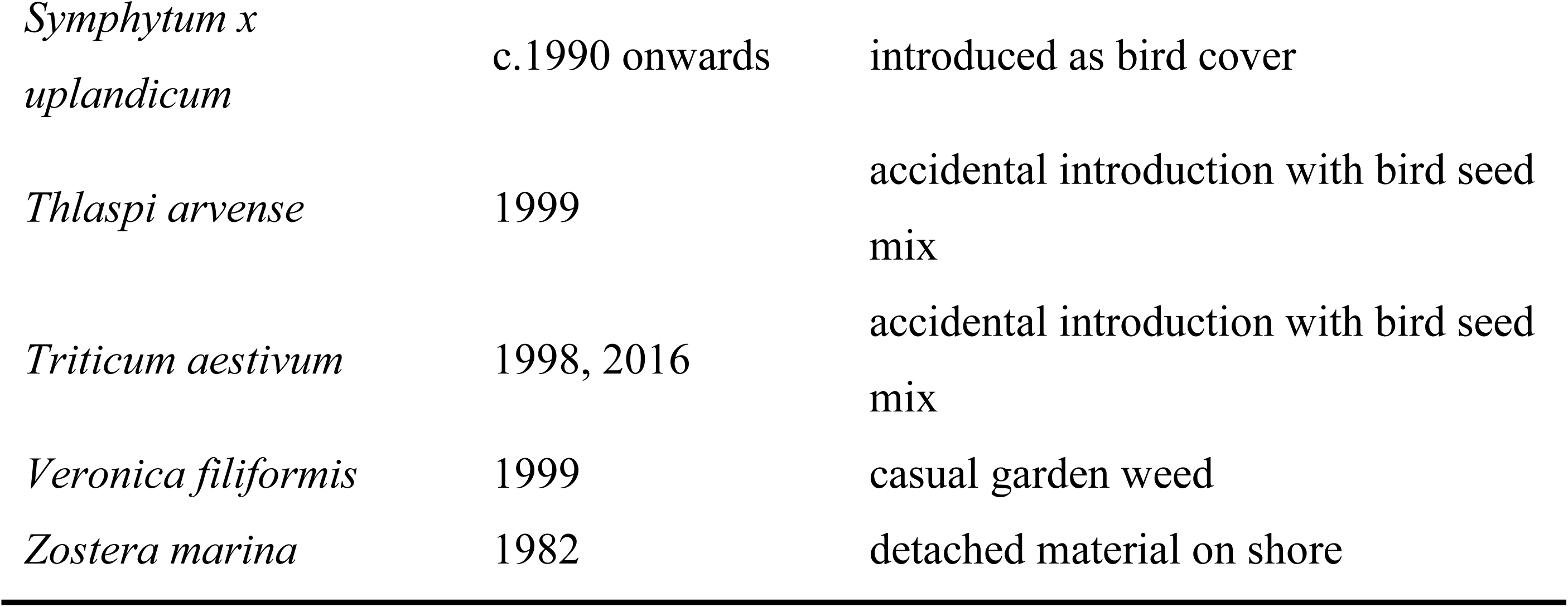
Additions to Fair Isle flora post-Scott & Palmer (1987). Those in bold were not reported for Shetland in Scott & Palmer (1987) and are thus cited for the first time in the archipelago. Likely source of origin is reported in the final column.

A number of species were found to be highly abundant across the island. The ten most widespread species were found in almost all survey squares. *Armeria maritima, Cerastium fontanum, Festuca rubra* and *Silene uniflora* were found in all the monads, and across an extremely diverse range of habitats. *Aira praecox, Carex nigra, Cochlearia danica, Holcus lanatus, Plantago coronopus* and *Plantago maritima* were found in 18 of 19 monads (95%). As well as being widespread, many of these species were extremely abundant in a given monad and appeared to play a disproportionately important role in these ecosystems.

A total of 10 species of conservation concern were recorded (Table 4). They comprised four categorised as vulnerable: *Chenopodium murale, Coeloglossum viride* (Figure 2A), *Gentianella campestris* (Figure 2B) and *Spergula arvensis*; five near threatened: *Hymenophyllum wilsonii* (Figure 2C), *Mertensia maritima* (Figure 2D), *Radiola linoides* (Figure 2E), *Viola tricolor* and *Zostera marina*, and one endangered: *Euphrasia marshallii*. According to the BSBI (2016), the species listed as nationally scarce on Fair Isle are: *Euphrasia foulaensis, Euphrasia ostenfeldii, Mertensia maritima, Odontites vernus, Ophioglossum azoricum, Polygonum boreale* and *Puccinellia distans* subsp. *borealis*. One species is listed as nationally rare, which is *Euphrasia marshallii*.

**Table 4.** Species diversity by monad for Fair Isle. All monads belong to national grid square HZ. For native status N: native, AR: archaeophytes, AN: neophyte. For Vascular Plant Red Data List for Great Britain NT: near threatened, VU: vulnerable. For Botanical Society of Britain & Ireland (BSBI) status NS: nationally scarce.

| N° | Monad<br>(HZ) | Species per<br>monad | Native status |  |  | Red Data List |  | BSBI<br>status |
| --- | --- | --- | --- | --- | --- | --- | --- | --- |
|  |  |  | N | AR | AN | NT | VU | NS |
| 1 | 1969 | 78 | 75 | 2 | 1 | 1 | 1 | 6 |
| 2 | 2069 | 117 | 111 | 4 | 2 | 1 |  | 4 |
| 3 | 2169 | 47 | 47 |  |  |  |  | 3 |
| 4 | 1970 | 92 | 90 | 1 | 1 | 1 |  | 2 |
| 5 | 2070 | 165 | 145 | 12 | 8 | 1 | 2 | 1 |
| 6 | 2170 | 86 | 86 |  |  |  |  | 2 |
| 7 | 2270 | 9 | 9 |  |  |  |  | 1 |
| 8 | 1971 | 29 | 29 |  |  |  |  | 1 |
| 9 | 2071 | 156 | 141 | 5 | 10 | 1 | 2 | 2 |
| 10 | 2171 | 152 | 144 | 2 | 6 | 2 | 1 | 4 |
| 11 | 2271 | 55 | 54 | 1 |  | 1 |  | 3 |
| 12 | 2072 | 103 | 103 |  |  | 1 | 1 | 3 |
| 13 | 2172 | 108 | 107 |  | 1 | 1 |  | 3 |
| 14 | 2272 | 130 | 116 | 4 | 10 | 1 | 1 | 4 |
| 15 | 2073 | 84 | 84 |  |  | 2 |  | 1 |
| 16 | 2173 | 114 | 114 |  |  | 1 | 1 | 2 |
| 17 | 2273 | 99 | 99 |  |  | 2 |  | 3 |
| 18 | 2074 | 21 | 21 |  |  |  |  | 1 |
| 19 | 2274 | 49 | 48 |  | 1 |  | 1 | 1 |

**Figure 2.**
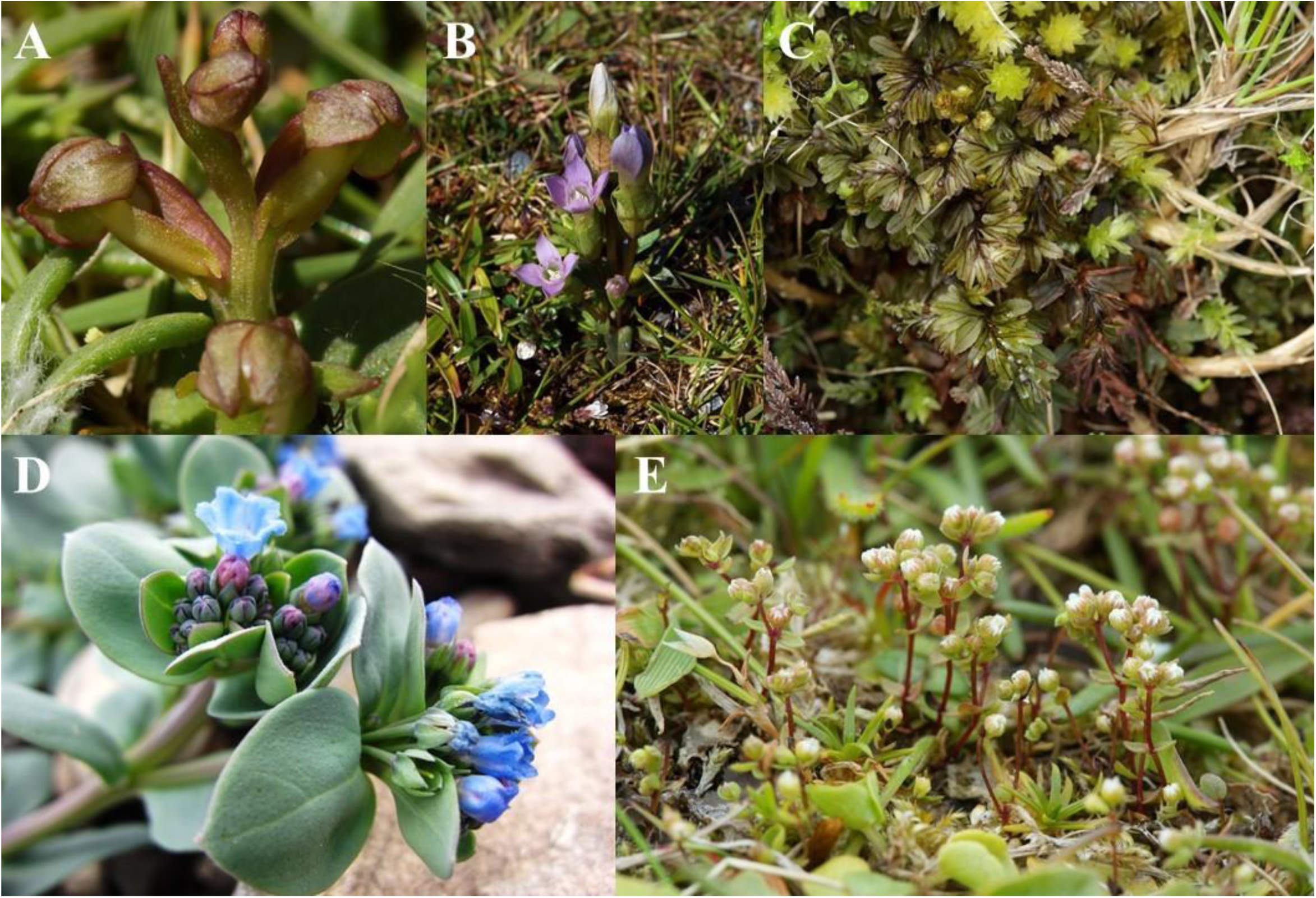
Threatened species present on Fair Isle as classified by the Vascular Plant Red List (Cheffings *et al*., 2005): (A) *Coeloglossum viride* (VU), (B) *Gentianella campestris* (VU), (C) *Hymenophyllum wilsonii* (NT), (D) *Mertensia maritima* (NT), (E) *Radiola linoides* (NT).

A total of 254 native species (80% of total island diversity) were recorded, with the remaining species split between archaeophytes (31 species) and neophytes (32). The most widespread alien taxa was *Matricaria discoidea* (present in nine monads), while nearly two thirds (21/33) of the other aliens were restricted to one or two monads.

### Habitat diversity

A total of 690 hectares out of 768 hectares were successfully surveyed for habitat type, the 78 remaining hectares that were not are allocated fall into the Phase 1 Miscellaneous category (Table 5), which incorporates developed areas such as active quarries, an airstrip and roads. Twelve major habitat types, plus 29 habitat sub-divisions (Figure 3), were found in the 1991-1992 Phase 1 Habitat Survey. In the follow-on habitat survey of June 2016 all the habitats from the Phase 1 survey were recognized with no large-scale changes.

**Table 5.** Main habitat categories as per Phase 1 Habitat Survey.

| Habitat | Percentage |
| --- | --- |
| Heathland | 31% |
| Coastland | 28% |
| Grassland | 19% |
| Miscellaneous | 10% |
| Mire, flush & spring | 4% |
| Arable | 3% |
| Marsh | 3% |
| Tall herb & fern | 1% |
| Rock exposure (inland cliff) & waste | 1% |
| Open water | <1% |
| Swamp & marginal | <1% |
| Woodland | <1% |
| **Total** | **100%** |

**Figure 3.**
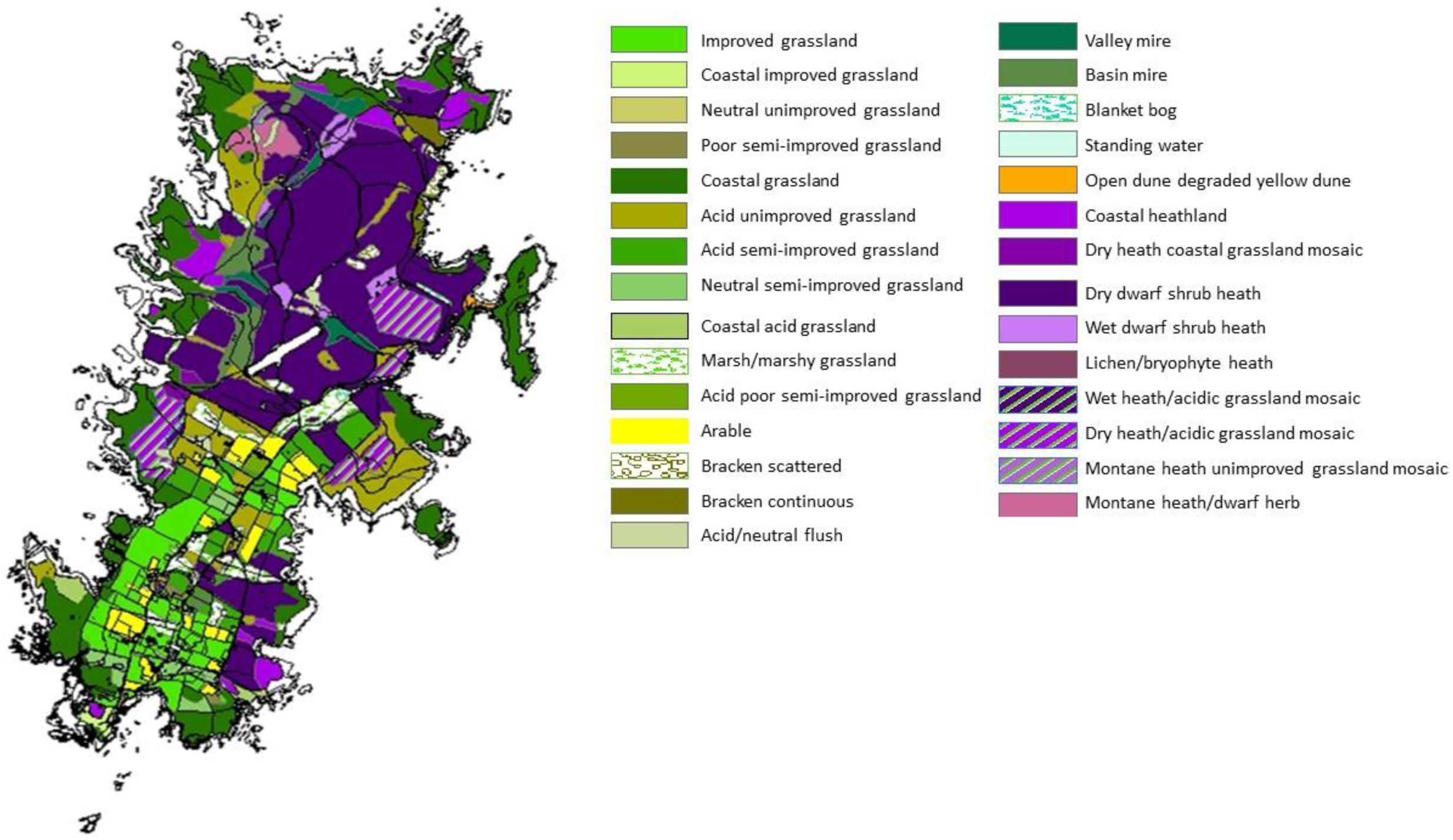
Map of vegetation of Fair Isle according to the results of the Phase 1 Habitat Survey.

## Discussion

This study aimed to provide the first comprehensive floristic analysis of the remote island of Fair Isle. We achieved this through generating a complete checklist of the flora, identifying and mapping species distributions and their relationships to habitats, and evaluating species considered under threat. Here, we discuss how the island’s vascular plant diversity has been assembled over time, and compare the diversity with other remote islands. We also suggest fruitful routes for future biological monitoring.

### Total species diversity on Fair Isle

Fair Isle has a total of 317 species, a subset of those from the total flora of Shetland (827 species, Scott & Palmer, 1987) and flora of Britain and Ireland (2951 species, Preston *et al*., 2002). Considering the land area of Fair Isle, which is just 7.68 km^2^, compared to Shetland which is 200 times the size at 1,466 km^2^, we can see that this value compares favourably. While a common assemblage of widespread species is expected on any landmass in Britain, this value is higher than many other remote islands, pointing to Fair Isle’s diverse range of habitats (discussed below). It is still notable, however, that 15 widespread species found commonly across Shetland have traditionally been thought to be absent from Fair Isle (Scott, 1972). These species are: *Equisetum fluviatile, Thalictrum alpinum, Drosera rotundifolia, Anthriscus sylvestris, Erica tetralix, Menyanthes trifoliata, Myosotis secunda, Veronica officinalis, Veronica serpyllifolia, Solidago virgaurea, Cirsium palustre, Luzula pilosa, Juncus effusus, Eleocharis multicaulis* and *Carex ovalis*. However, we have now found that three of these species (*Drosera rotundifolia, Juncus effusus* and *Carex ovalis* (= *C. leporina*)) are present. In addition to a diversity of higher plants, the island is emerging as a diverse area for other species groups, for instance 170 bryophyte species including strong populations of the nationally rare *Sanionia orthothecioides* (Riddiford, Unpublished; S. Payne, pers. comm.), 294 lichen taxa and 13 lichenicolous fungi including 14 classified as nationally rare and 55 as nationally scarce (Price, 2017) and a range of nationally scarce invertebrates including two species new to the British Isles (Riddiford & Young, 2016; Disney & Riddiford, 2017), a post-ice age relict Heteropteran (Riddiford, 2010) and terrestrial and marine species close to the northern or southern boundaries of their range (Riddiford & Riddiford, 2011; Riddiford & Shaw, 2011; Riddiford, 2016).

The Fair Isle flora includes representatives from 31 orders, 68 families and 191 genera. Three families, Asteraceae (37 species), Poaceae (37 species) and Cyperaceae (20 species) contributed disproportionately to the total island diversity. These groups are likely to be well represented for a number of reasons. Firstly, these groups are often the most abundant in temperate floras and as such should be expected to be common. Secondly, Asteraceae and Poales possess wind dispersed seeds and fruits (Simpson, 2006), allowing them to colonise the remote island. Thirdly, traditional crofting maintains grassland across the island, thus there is plenty of suitable habitat. Finally, the prevalence of asexual reproduction (*e.g*. rhizomes, bulbils, stolons) may allow these groups to colonise despite lack of conspecific mates with which to reproduce.

While overall species diversity appears to be relatively high, our survey revealed a scarcity of alien taxa on the island. Over 80% of the flora of Fair Isle are native species, much higher than the 66% for Shetland (Scott & Palmer, 1987) and the 52% for the UK (Preston *et al*., 2002). The paucity of alien species could be due to both potential for dispersal, and for establishment. In terms of dispersal, the isolated nature of Fair Isle may limit opportunities for plant species that have recently been introduced to the UK to naturally disperse to the island. This is supported by most alien taxa on Fair Isle being of horticultural value, rather than representing widely dispersing non-horticultural taxa (the exceptions here being the dispersive *Matricaria discoidea* and *Capsella bursa-pastoris*). In terms of establishment, the harsh environment on Fair Isle will prevent the persistence of propagules poorly adapted to these novel conditions.

### Conservation of species diversity

Ten species on Fair Isle are included in the Vascular Plant Red Data List for Great Britain (Cheffings *et al*., 2005): four are listed as vulnerable (*Chenopodium murale, Coeloglossum viride, Gentianella campestris* and *Spergula arvensis*), five as near threatened (*Hymenophyllum wilsonii, Mertensia maritima, Radiola linoides, Viola tricolor* and *Zostera marina*) and one, *Euphrasia marshallii*, as endangered.

Some of these taxa are relatively widespread on Fair Isle but are undergoing rapid decline across Great Britain. *Coeloglossum viride, Gentianella campestris* and *Spergula arvensis* (all vulnerable) have been recorded on Fair Isle since Trail (1906), indicating their presence on the island for at least a century. Species classified as “near threatened” have been added to the island list successively with additional recording effort (*e.g*. *Hymenophyllum wilsonii*, Hannah Stout in Fitter (1959); *Radiola linoides*, Currie (1960); *Mertensia maritima*, N J Riddiford & P V Harvey in 1992). Given its scarcity on the island, *H. wilsonii* is more likely to have been overlooked than to be a recent colonist and *R. linoides*, which occurs across the island in very large numbers, may have been overlooked because of its diminutive size; previous colonisation by *M. maritima* may have been stymied by its vulnerability to livestock grazing.

Seven species, *Euphrasia foulaensis, Euphrasia ostenfeldii, Mertensia maritima, Odontites vernus, Ophioglossum azoricum, Polygonum boreale* and *Puccinellia distans* subsp. *borealis* are listed as nationally scarce (BSBI, 2016). In many cases, it is the nationally scarce species, which are recorded in 16-100 hectads (10km2), which are of great interest ecologically, as they are indicators of good quality habitat (Lockton *et al*., 2005). Many of these nationally scarce species have been declining due to habitat destruction over the last 40 years, and the stability of habitats and relatively unchanged land-use on Fair Isle may preserve scarce species that are declining elsewhere.

One species, *Euphrasia marshallii*, is nationally rare, supposedly found in 15 or fewer hectads in Great Britain (Lockton *et al*., 2005). This species is challenging to identify in the field and is morphologically similar to *E. ostenfeldii*, a species with which it frequently coexists. The following *Euphrasia* species have been published for Fair Isle (dates refer to year of publication (*i.e*. Scott, 1972; Scott & Palmer, 1987; Scott, 2011)): *Euphrasia nemorosa* (1987), *Euphrasia confusa* (1972, 1987), *Euphrasia foulaensis* (1972, 1987), *Euphrasia ostenfeldii* (1987), *Euphrasia micrantha* (1972 (det. P. F. Yeo), 1987), *Euphrasia scottica* (1972), *Euphrasia foulaensis x marshallii* (1972), *Euphrasia marshallii* (1972), *Euphrasia confusa x foulaensis* (1972, ‘best omitted as not positively identified’ (Scott & Palmer, 1987)), *Euphrasia arctica* (reported as *E. borealis*, 1972). However, the taxonomic status of *Euphrasia* on Fair Isle, and indeed *Euphrasia* taxonomy across the UK, is still uncertain. UK-wide *Euphrasia* surveys, in conjunction with genetic analysis, are needed to reveal the true status of these complex tetraploid species in the UK. Other surveying work on Fair Isle should be done for taxonomically challenging *Carex, Agrostis, Deschampsia* and *Atriplex*, which remain uncertain.

The survey method employed was not chosen to look at changes in species abundance. However, islanders have provided verbal accounts of changes which would be worthy of future investigation. Though speculative, it is likely that climate change may affect species distributions and this is likely to be of great importance in the future. This is particularly the case for species with small population sizes and at the edge of their climatic range (*e.g. Radiola linoides, Apium inundatum* and *Hymenophyllum wilsonii*).

### Habitat diversity on Fair Isle and its relationship with species diversity

Twelve habitats and 29 habitat sub-divisions were recorded on the island, demonstrating considerable habitat variation for a small area. Fair Isle’s open landscape and small size permitted full access to the area and comprehensive assessment of nearly every habitat. Whereas some of the habitats (*e.g*. dry dwarf shrub heath) were extensive, Fair Isle has a considerable number of ‘niche’ habitats which are small but significant in terms of their communities. Comparisons between habitat surveys (1991-1992; 2016) revealed most areas had not experienced major changes in land use or habitat type. The only exception is arable land, where in the 1991-92 survey there were 12 arable rigs (meaning a croft with a rig of oat, potato or turnip crops, plus a general vegetable plot). In 2016, there were six arable rigs. The rigs were mainly alongside grass leys cut for hay or silage and have now been incorporated into the silage fields. The others fall into the improved grassland category. In the 1991-92 survey, NCC Phase 1 categories were followed so the 1991-92 map incorporated rigs and grass leys as Arable without differentiation.

Despite Fair Isle’s diversity of habitats, it is notable that these are only composed of low-growing or shrubby plants. Even casual observers will notice that Fair Isle is virtually treeless, apart from a few species introduced by the Fair Isle Bird Observatory in enclosed sites. This reflects the intense grazing pressure the island has been subjected to, be it from ponies and cattle (pre-20^th^ century), or more recently from sheep. There is historical evidence to suggest juniper and *Salix* may have once been more widespread, and a single survivor of *Salix cinerea* subsp. *oleifolia* persists on the island (at the coastal end of Funniequoy Gully). Grazing from sheep, even in apparently inaccessible areas, has further reduced plant height and seems to be preventing habitats reaching a potential climax vegetation of low scrub composed of limited but rich diversity. Even if grazing was prevented, for example by enclosures, it seems unlikely that succession would give rise to woodland. Shetland in general is depauperate of trees, and the extreme environment seems to select against tall woody vegetation.

Attempts to classify and describe Fair Isle plant communities using the National Vegetation Classification (NVC) have been made by O’Hanrahan (2003) and Quinteros (2016). Both studies showed several species occur across a wide range of communities, resulting in significant deviations from standard NVC communities. This may in part be caused by the effect of the extreme oceanic climatic regime on the floristic composition of the communities, as well as the high degree of habitat mixing resulting in species ‘spilling over’ into adjacent habitats.

Internationally, the British Isles are known to have a large expanse of heather-dominated vegetation of mostly Ericacae (*e.g*. *Calluna vulgaris, Empetrum nigrum* and *Erica cinerea*). Fair Isle has large areas of such habitat with, in addition, a strong population of the prostrate form of juniper *Juniperus communis* subsp. *nana* which is otherwise very sparse in Shetland (O’Hanrahan, 2003). The importance of this habitat for birds is recognised in its designation as a Special Protection Area (SPA) under the EU Birds Directive; and in providing protection for smaller plants and non-vascular plants in its designation as a Special Area of Conservation (SAC) for its dwarf shrub heath and maritime habitats under the EU Habitats Directive. Research on the main types of biological and climatic damage associated with this habitat is suggested by this paper.

### Diversity on Fair Isle compared with other islands

There are a lot of factors that complicate the comparison of flora between islands, including geology, history, isolation, distance from the mainland, climate, human impact as well as factors affecting genetic variation and local adaptation. Even so, some general observations can be made. The flora of Fair Isle is comparable in many respects with the remote Scottish islands of St Kilda (Crawley, 2014) and Foula (Barkham *et al*., 1981; BSBI, 2015), which we use here for comparison.

St Kilda is an isolated island 64 km (40 miles) off the west coast of Scotland, with a land area of 854 ha. The flora of St Kilda has been studied extensively for over 20 years, with the most recent update made in 2014. St Kilda is relatively depauperate with just 195 species (Crawley, 2014). Both islands share a considerable number of species, with 79% of species on St Kilda found on Fair Isle. Thus, there appears to be a common pool of widespread species shared between two of Britain’s most isolated islands, with differences likely due to chance colonisation and establishment. Another interesting feature of St Kilda is that all but two species (*i.e*. *Matricaria discoidea* and *Epilobium brunnescens*) found on the island are native, thus it supports fewer alien species than Fair Isle, perhaps due to less movement of people from the mainland.

Foula is one of Great Britain’s most remote permanently inhabited islands. It is located 32 km west of Walls in Shetland, and has an area nearly twice the size of Fair Isle, at 1,265 ha. According to Barkham *et al*. (1981) and the BSBI (2015), Foula has an estimated 316 plant species. Of the total number of species present on Foula, 66% are present on Fair Isle. This again highlights the stochastic nature of colonisation associated with remote islands: that while both islands share many species, there are a substantial number which have not successfully colonised both islands.

### Conservation recommendations for Fair Isle

An up to date flora, with an associated overview of the plant communities, is a good launching pad for future botanical research and conservation efforts. Fair Isle has a number of species on the Vascular Plant Red List. These species must be evaluated in terms of their habitat requirements before decisions are made on a programme of conservation, in order to establish whether a species-based or habitat-based action plan is required. As a first step, a monitoring programme is imperative, linked to a management plan that can act in response to future climate change and potential changes in land-use.

For example, it is important to understand population trends for montane, arctic-alpine and relict species (*e.g*. *Salix herbacea, Polygonum viviparum, Carex bigelowii, Trientalis europaea*) at a time of climate change. Sensitive arctic–alpine species are found at unusually low elevations on Fair Isle (just 207 meters’ elevation) and could serve as an early warning indicator of the effects of climate change. This is particularly relevant as detectable changes in weather patterns have already been observed on the island, including higher average temperatures and wetter winters since the 1970s (Wood and Bunce, 2016). An advantage of monitoring Fair Isle, rather than other areas, is that its small size and defined habitats makes it an easy study area. Its suitability is further complemented by access to a 40+ years run of weather data available through the Fair Isle Meteorological Station.

The Red List for the British Isles (Cheffings *et al*., 2005) discusses plants that are remarkably difficult to include in threat assessments. Much of the difficulty associated with these groups comes from species identification problems. Fair Isle is a hotspot for two groups: *Euphrasia*, where eight species were identified during this survey, and *Dactylorhiza*, where two species were noted. *Euphrasia* is a highly critical genus with over 60 wild hybrids and active speciation (Cheffings *et al*. 2005), and several of the emergent taxa are narrowly distributed and endemic to the British Isles. *Dactylorhiza* is a difficult genus owing to hybridisation between almost all the taxa and active speciation is evidently taking place (Cheffings *et al*. 2005). Work with specialist taxonomists is suggested for future surveys.

In terms of planning future habitat management, it will be important to undertake detailed surveys of bracken (*Pteridium aquilinum*). This species is found in five monads of heathland on the north side of the island, and has been present for at least 100 years (the late James A. Stout, pers. comm.) Bracken spreads by underground rhizomes and prevents other species growing (Proctor, 2013). It was always very limited in distribution, but expanded in the 1980s, both in density and range. The extent to which it competes with *Calluna vulgaris* is unclear and it would be important to observe future distribution changes.

In November 2016, Fair Isle and its surrounding waters became Scotland’s first Demonstration and Research Marine Protected Area (MPA). The scheme is being taken forward by the Fair Isle community in partnership with other stakeholders, in particular the fishing industry and the scientific body. The community had petitioned the Scottish Government for an MPA over a considerable number of years, driven by observations of marked negative changes in the maritime resource. Designation provides a platform for widening the knowledge base which in turn guides sustainable management of that resource. It is already apparent from several long-term Fair Isle datasets that climate is a major player in the changes observed by the islanders. Aspects of Fair Isle flora have a role in the overall study of climate-driven change, for instance impacts on: arctic-alpine species such as *Salix herbacea*, restricted to Fair Isle highest points on the Fair Isle; the *Mertensia maritima* population in the face of increasing frequency and strength of storms; and population gain or loss for hyperoceanic species such as *Ophioglossum azoricum* (and other biota amongst the bryophytes and lichens). This places Fair Isle flora in its maritime context and the potential to provide added value to the monitoring programme within the MPA research framework and goals.

## Supplementary information

Supplementary Text S1. Habitat description and histories for each of the 19 monads surveyed.

Supplementary Table S2. Categories for Fair Isle status.

## Acknowledgements

We are very grateful to Sally Eaton for advice and supervision; Jenny Farrar & Jo Whatmough for their guidance in fieldwork and plant identification; The Fair Isle Bird Observatory Trust and its wardens, Susannah and David Parnaby, for use of facilities at the Bird Observatory; Lee Gregory for his help in the field and photographic material; Steven Sylvester for grasses identification; and Jim McIntosh for project guidance and providing access to the BSBI Distribution Database. This study formed part of CQP’s MSc Thesis at the Royal Botanic Garden Edinburgh (RBGE), supported by the Scottish Government’s Rural and Environmental Science and Analytical Services Division. Additional funds were provided by the RBGE Sibbald Trust. Research by ADT is supported by NERC Fellowship NE/L011336/1. This paper is dedicated to the memory of Tessa McGregor, a field biologist who encouraged this research.

